# Targeted curation of the gut microbial gene content modulating human cardiovascular disease

**DOI:** 10.1101/2022.06.20.496735

**Authors:** Mikayla A. Borton, Michael Shaffer, David W. Hoyt, Ruisheng Jiang, Jared Ellenbogen, Samuel Purvine, Carrie D. Nicora, Elizabeth K. Eder, Allison R. Wong, A. George Smulian, Mary S. Lipton, Joseph A. Krzycki, Kelly C. Wrighton

## Abstract

Despite the promise of the gut microbiome to forecast human health, few studies expose the microbial functions underpinning such predictions. To comprehensively inventory gut microorganisms and their gene content that control trimethylamine induced cardiovascular disease, we mined over 200,000 gut-derived genomes from cultivated and uncultivated microbial lineages. Creating MAGICdb (Methylated Amine Gene Inventory of Catabolism database), we designated an atherosclerotic profile for 6,341 microbial genomes that encoded metabolisms associated with heart disease. We used MAGICdb to evaluate diverse human fecal metatranscriptome and metaproteome datasets, demonstrating how this resource eases the recovery of methylated amine gene content previously obscured in microbiome datasets. From the feces of healthy and diseased subjects, we show MAGICdb gene markers predicted cardiovascular disease as effectively as traditional blood diagnostics. This functional microbiome catalog is a public, exploitable resource, enabling a new era of microbiota-based therapeutics.

## Introduction

Mounting evidence implicates the gut microbiome, the thousands of microorganisms and their gene products residing in the gut, as a critical modulator of human health^1,2^. One of the most compelling examples implicating gut microbial metabolism as a factor in human disease is atherosclerotic cardiovascular disease (ACVD), which is the leading cause of death globally^3–5^. Here gut microorganisms process quaternary amines from protein rich foods (e.g. eggs, beans, meat) to generate the metabolite trimethylamine (TMA, **Figure 1)**. TMA, an obligate microbiota derived metabolite, is absorbed into the blood stream and subsequently transformed by the liver to trimethylamine-N-oxide, which promotes ACVD in humans^3,5^. Gut microorganisms also catalyze reactions that reduce gut TMA concentrations^6^. While linkages between the gut microbiota and atherosclerosis are accepted^3–5,7^, a systems microbiology approach to reveal the balance between these reactions and their contributions to TMA production has yet to be holistically applied in the human gut.

**Figure 1.**
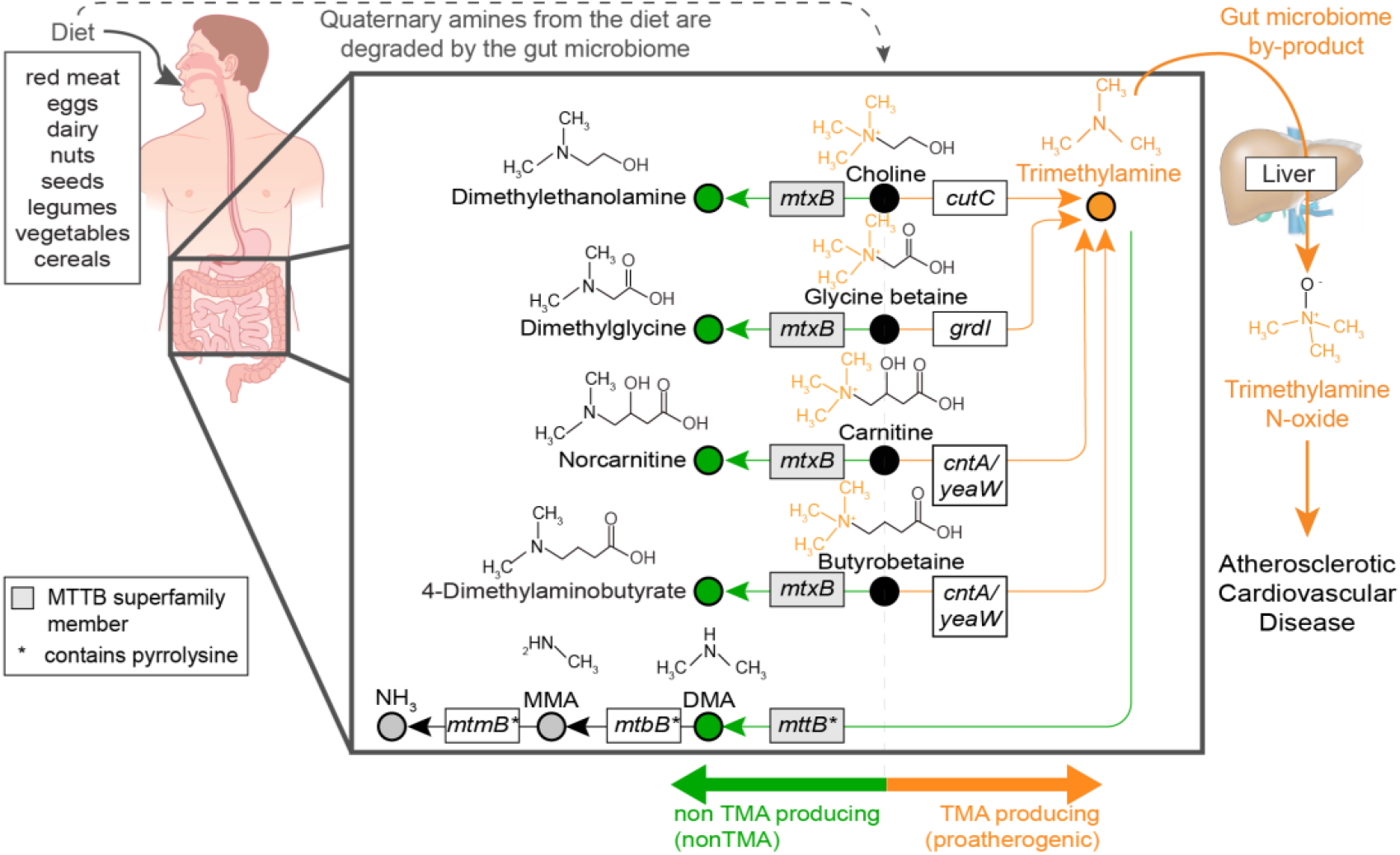
Microbial proatherogenic or nonatherogenic methylated amine metabolic routes for dietary quaternary amine transformations in the gut. Foods in the human diet, including read meat and certain vegetables, have elevated quaternary amines^71^. Upon consumption, these compounds travel to the gut where they are degraded by microorganisms. Chemical structures for dietary quaternary amines (black circles) are shown for choline, glycine betaine, carnitine, and butyrobetaine, with the trimethylamine (TMA) moiety of these compounds noted in orange. Microbial proatherogenic conversions (orange arrow) of these compounds yield TMA (orange circle), which is exported to the liver where human enzymes convert TMA to Trimethylamine-N-oxide, a metabolite that promotes atherosclerosis. Alternatively, microorganisms catalyze demethylation reactions that subvert TMA concentrations (green arrows). In the first route, microbial dietary quaternary amine processing does not result in TMA production, but instead yields non-TMA metabolites (green circles) like dimethylethanolamine, dimethylglycine, norcarnitine, or 4-dimethylaminobutyrate. In a second TMA-reducing route, TMA is directly microbially demethylated to dimethylamine (DMA, green circle). Sequential demethylations of DMA to MMA (monomethylamine) and MMA to ammonium are also noted by grey circles. For each conversion, the microbial abbreviated gene names are noted in grey or white boxes, with the full gene names, reactions, and citations included in **Table S1**.

The microbial biochemistry catalyzing TMA production from quaternary amines in the gut are commonly attributed to four routes from dietary choline^8^, glycine betaine^9^, carnitine^10^ and butyrobetaine^11^ (**Figure 1**, orange). Alternatively, through demethylation reactions, microorganisms can either directly reduce TMA concentrations^12^ or act on dietary quaternary amines^6,13–17^ to indirectly subvert TMA production (**Figure 1**, green). These nonTMA producing demethylation reactions are catalyzed by enzymes belonging to the same superfamily (MTTB), with the TMA specific enzymes distinguished by the presence of a pyrrolysine amino acid^6,18,19^. Despite their capacity to reduce concentrations of disease-causing TMA, the nonTMA routes remain largely enigmatic in the human gut today.

Collectively the microbial conversions of quaternary amines and their derivatives are here referred to as methylated amine (MA) metabolism. MA metabolism has been poorly characterized in the gut for several reasons. First, many of the nonTMA producing, quaternary amine acting enzymes were discovered in the past five years^13–16^, and are sampled only from a few cultivated microorganisms (**Table S1**). Additionally, both proatherogenic and nonTMA producing MA enzymes are either misannotated or not annotated at all in automated workflows commonly used to characterize the gut microbiome. Complications include misannotations due to close homology with genes of different functions that then require manual active site confirmation (e.g. *cutC, grdI*)^20,21^. Also, within the MTTB superfamily many members either have unknown biochemical functions (e.g. *mtxB*)^6^ or the presence of pyrrolysine results in truncated genes (e.g. *mttB*) (**Figure 1**)^12,18,19^. Beyond annotation, many of the known TMA-utilizing microorganisms, like methanogens, are rare members in the gut and are often missed because of sampling considerations^22,23^. These challenges collectively result in an incomplete understanding of this important, disease relevant gut metabolism.

Given their potential contributions to atherosclerotic cardiovascular disease (ACVD), we hypothesized the complete profiling of the proatherogenic (TMA producing) and nonTMA producing genes in fecal content would discriminate healthy and diseased individuals. To test this hypothesis, we cataloged the proatherogenic and nonTMA producing gene content from more than 200,000 microbial genomes derived from the human gut^24–26^. We constructed the Methylated Amine Gene Inventory of Catabolism database (MAGICdb), a resource that uncovered the untapped atherosclerotic disease modulating microorganisms prevalent and metabolically active in the human gut. We then used MAGICdb to diagnose atherosclerotic cardiovascular disease from fecal microbiomes, demonstrating a diagnostic performance on par with traditional lipid blood markers. This microbial genome resource paves the way for disease diagnoses and management from microbiome content, representing a new avenue for the development of therapeutic interventions in precision medicine.

### Methylated amine transformations are a keystone metabolism in the human gut

To inventory the MA content in the gut microbiome, we developed a computational workflow that overcame prior annotation challenges by employing homology and non-homology approaches (**Figure S1**). We first identified homologs within each of the 7 gene types and then followed two distinct curation paths: one for the nonatherogenic superfamily members (*mtxB, mttB*) and other demethylating genes (*mtmB, mtbB*) and another for proatherogenic members (*cutC, cntA*/*yeaW, grdI*). Following manual curation of these genes, the microbial genomes were defined as proatherogenic (TMA producing), non-TMA producing, or both based on their collective gene content.

To link fecal TMA concentrations to microbial gene content we performed a human cohort study of 113 individuals (**Figure 2A, Figure S2AB**). We applied our computational workflow to 54 fecal metagenomes that spanned quartiles of fecal TMA concentrations derived from our cohort (**Figure 2B**). To sample rare microbial members, these fecal metagenomes were sequenced up to 55 Gbp/sample (mean of 18 Gbp/sample, **Table S1**), resulting in deeper sequencing than the 4 Gbp/sample that is commonly used in gut metagenome studies^27–29^. We show that a metagenome sequence depth of more than 35Gbp recovered nearly double the amount of MA genes than the traditional 4 Gbp (**Figure S3**). At a cumulative sequencing depth of 775 Gbp (i.e., 75% of our total sequencing) the MA gene discovery rate plateaued (**Figure 2C**). Beyond sequencing depth, gene recovery was also enhanced by the number of individuals sampled, suggesting this metabolism may be variably dispersed across humans (**Figure 2DE**). These analyses illuminate the importance of considering gene abundance and cohort distribution when designing experiments to target specific metabolisms from a complex microbiome like the gut.

**Figure 2.**
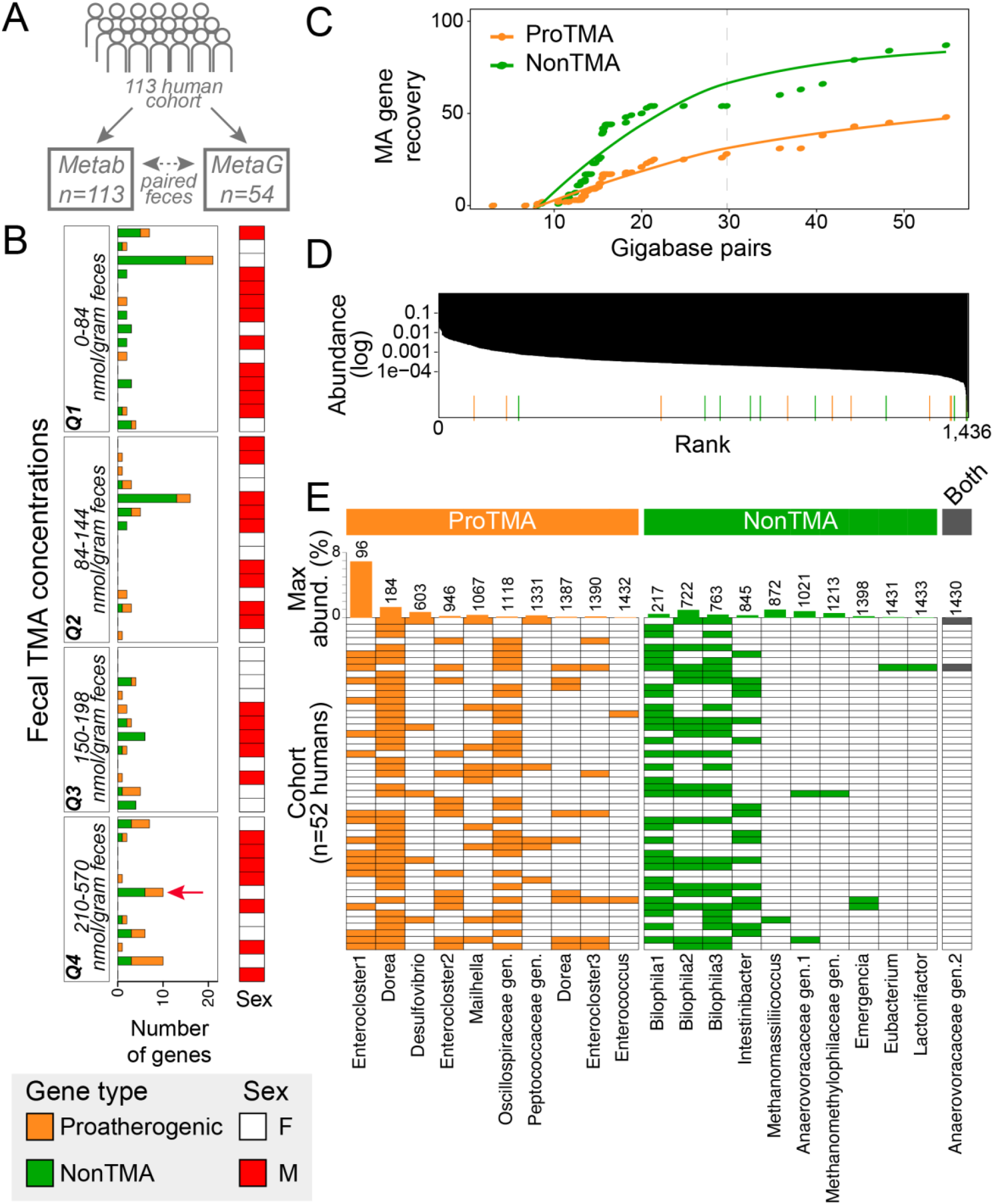
Microbially methylated amine utilization is encoded by rare members with differential occupancy sampled across a human cohort. **A** A 113 human cohort study resulted in metabolite analysis (Metab) of fecal TMA concentrations, which were assigned to quartiles (Q1-Q4) based on concentration. Using these quartiles, 54 samples were selected for metagenomics (MetaG), with at least 12 samples chosen from each quartile. **B** Quantification of the proatherogenic (orange) and nonTMA (green) genes inventoried in each fecal metagenome, organized by quartile, with quartile ranges noted in the left panel and sex of the subject noted in the right panel. Red arrow denotes the fecal sample chosen for quaternary amine amendment in Figure 4. **C** MA discovery curve denotes the number of new genes recovered with increased sequencing depth. The dashed line indicates the plateau of new MA gene recovery. **D** A rank abundance curve of the average relative abundance of 1,436 MAGs (y axis) and their average rank (x-axis) in each sample sequenced as part of this cohort. The average ranked relative abundance of MA containing genomes are highlighted by colored bars along the x-axis. **E** The presence (filled) and absence (white) of MA containing MAGs, with the bar graphs at the top reporting the maximum relative abundance and average rank (numbers out of 1,436) of each genome. In B-E, colors correspond to proatherogenic (orange), nonTMA producing (green), or both (black), based on MA content as defined in **Figure 1**.

From our cohort we sampled 153 MA genes (135 unique) with 41% and 59% of the genes given proatherogenic or nonTMA assignments respectively (**Table S1**). We found no considerable relationship between gene content or TMA concentrations with host sex, body mass index, or lifestyle (**Figure S2)**. As expected from relationships outlined in **Figure 1**, the relative abundance of the proatherogenic *cntA*/*yeaW* gene was associated with higher fecal TMA concentrations, while the TMA reducing *mttB* and *mtxB* genes relative abundance was associated with lower TMA concentrations (**Figure S3B**). More importantly, while neither the proatherogenic nor the nonTMA summed relative abundances were on their own able to predict fecal TMA concentrations, together their cumulative profile was predictive (**Figure S3D**). While deduced from a small sized cohort, this early data supports the notion that the comprehensive MA functional content previously obscured in feces may have explanatory relevance for the atherosclerotic status in humans.

Across the cohort we reconstructed 2,447 high and medium quality microbial metagenome assembled genomes (MAGs) that were dereplicated into 1,436 genomic representatives (**Table S1**). Only 1.5% of these dereplicated MAGs encoded at least one MA gene (**Figure 2D**). These 21 MA containing genomes were nearly equally classified as proatherogenic and nonTMA producing. Only a single genome, a member of a novel genus in the Anaerovoracaceae, had both TMA-forming and depleting genes. Confirming our earlier suspicions, microbial members that encoded MA genes, especially the less studied nonTMA types, were some of the rarest members in the fecal community and were heterogeneously distributed across the cohort (**Figure 2DE**).

MAGs did not have consistent abundances or distributions across the cohort. The most dominant MA encoding genome was a proatherogenic *Enterocloster*. As only the 96th most abundant of the 1,436 MAGs sampled, this genome was detected in a third of individuals (**Figure 2E**). The second most abundant member, a proatherogenic *Dorea*, was sampled in nearly every human. Relative to their TMA producing counterparts, the four most dominant nonTMA producing genomes were less abundant but had similar occupancy across the cohort, with the remaining rare nonTMA members being more variably sampled (**Figure 2E**). Despite being encoded by rare and variably distributed members, the cumulative MA gene relative abundance was predictive of fecal TMA concentrations (**Figure S2**). Here we propose extending the idea of a microbial keystone species^30,31^ to the level of metabolism, establishing that in spite of low relative abundance methylated amine metabolisms in the gut have a disproportionately large effect on host physiology.

### Curation of methylated amine metabolism from over 200,000 microbial genomes

To create the most robustly sampled MA genome and gene resource, we extended sampling to ∼9,000 publicly available microbial fecal metagenomes^24,25^. This vastly expanded sampling beyond our cohort to thousands of humans spanning diverse ages, geographies, diets, and health conditions. We employed our computational workflow on 237,273 bacterial and archaeal MAGs. We also mined 700 bacterial genomes from gut microorganisms cultivated as part of the Human Microbiome Project^26^. In total, we analyzed the MA genes from 238,530 microbial genomes acquired from cultivated and uncultivated microorganisms (**Figure 3AB**). Showing the value of each of these datasets the large-scale MAG compendiums provided the most MA containing genomes, while our cohort-study derived MAGs and the microbial genomes from cultivated representatives provided a larger percentage of higher quality genomes that were maintained in the dereplicated database (**Figure 3B**).

**Figure 3.**
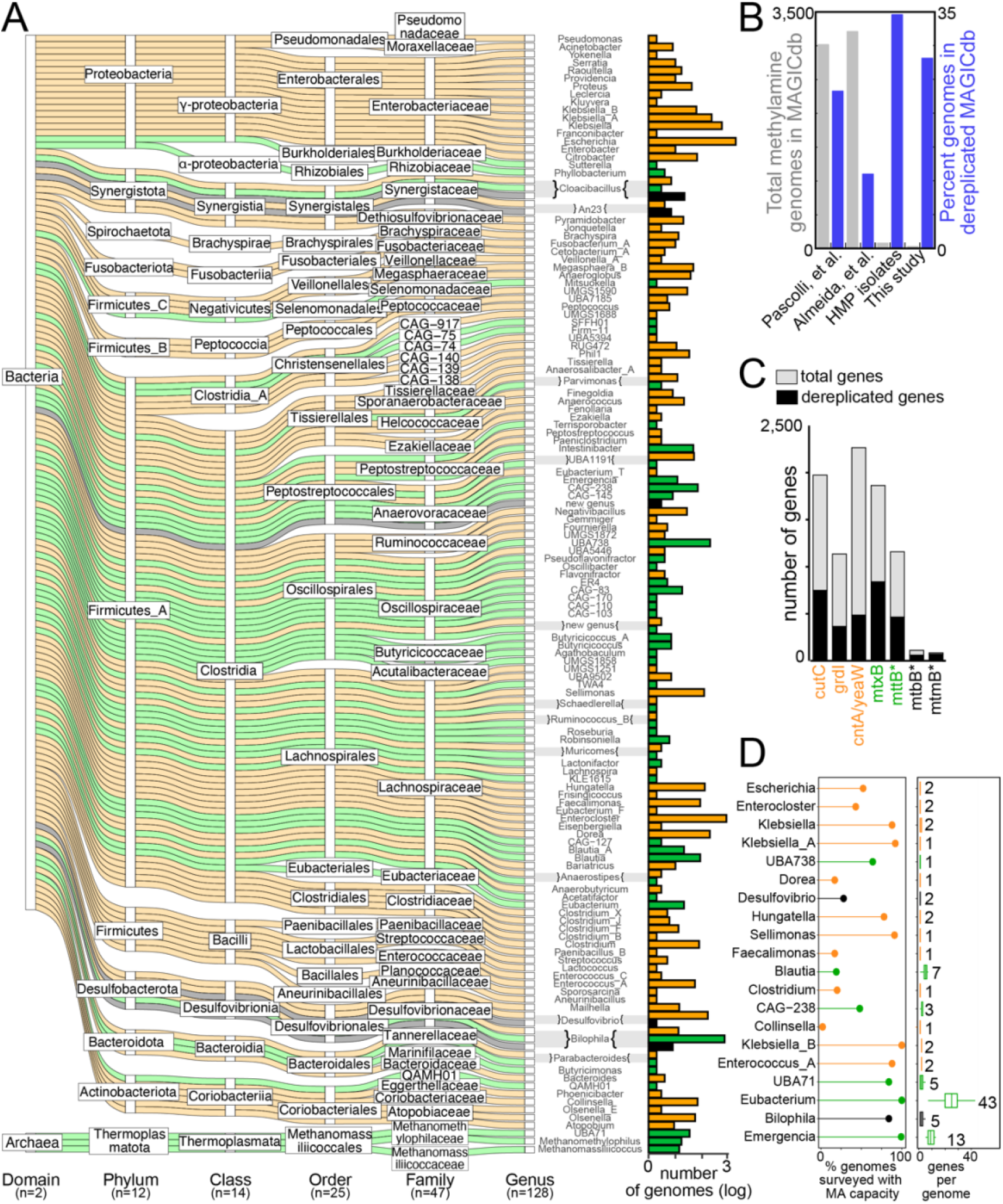
MAGICdb indexes the TMA relevant gene and genome content in the human gut microbiome. Throughout this figure TMA classifications are based on MA gene or genomic content as outlined in **Figure 1**, with colors denoting TMA status as proatherogenic (orange), nonTMA producing (green), or both (black). **A** Alluvial plot shows the taxonomic assignment of the 6,341 genomes that encode MA potential in MAGICdb. Alluvia are colored by MA genome content and TMA classification noted by coloring. The total number of genomes and their TMA classification(s) are summarized for each genus as a barchart. **B** The origin of the 6,341 genomes in MAGICdb (gray bars) and the percent of genomes that remained in MAGICdb after dereplication (blue bars). **C** At the gene level, a stacked bar chart reports the total (gray) and dereplicated (black) genes in MAGICdb, with asterisk indicating genes with a pyrrolysine amino acid. **D** The top 20 genera represented in MAGICdb and their TMA classification. For the genera with the most genomes sampled in the MAGICdb, the dot plot shows the percent of genomes surveyed within a genus with the capacity for MA metabolism, while the boxplots indicate the mean number and range of MA genes per genome within a genus. For each genus, the maximum number of MA genes in a genome is reported.

Mining this genome content, we created Methylated Amine Gene Inventory of Catabolism database (MAGICdb) which included (i) a gene dataset with unprecedented sampling of these disease relevant genes, and (ii) the corresponding linked genome dataset offering organismal context for MA metabolisms. MAGICdb contains 6,341 genomes encoding 8,721 MA genes (**Figure 3A, Table S2**). Within the MAGIC gene database, the proatherogenic and nonTMA gene richness was nearly equivalent (1,597 and 1,434 respectively) with *cutC* (choline trimethylamine lyase) and *mtxB* (non-pyrrolysine methyltransferase) being the most dominant types sampled respectively (**Figure 3C**). Considering the unique genes only, MAGICdb sampled up to 12-fold more genes compared to prior reports^20,22,32–35^. This expansion of MA gene diversity was attributed to the vast number of genomes collected from nonreference-based gut metagenome samples, rather than only relying on genomes from cultivated microorganisms like most prior analyses.

Within the MAGIC genome database, MA encoding members belonged to 1 archaeal and 11 bacterial phyla, or half of the phylum-level lineages surveyed (**Figure 3A**). This metabolism was found in less than 3% of the microbial gut genomes surveyed, indicating that even when scaled to a larger dataset this is a specialized metabolic capacity in the gut microbiome. Here we discovered the first MA containing genomes within the phyla *Spirochaetota* (12 MAGs exclusively proatherogenic) and *Synergistota* (59 MAGs with proatherogenic and nonTMA producing members). We also extended this metabolism to 88 gut genera that prior to our survey, were not recognized as playing a role in gut MA transformations (**Table S2**). This novelty sampled in MAGICdb documents the disease causing or ameliorating gene reservoir that was previously untapped within the gut microbiome.

Analysis of the TMA classifications across taxonomic levels revealed that all the Archaeal genomes were nonTMA producing. The same cohesive phylogenetic clustering of MA functionality was not observed for the bacteria at higher levels like class but was observed at finer taxonomic levels like genera. In fact, 89% of the 125 bacterial genera were exclusively proatherogenic (*Enterocloster, Citrobacter, Escherichia*) or nonTMA producing (*Eubacterium, Blautia*) (**Figure 3A**). The remaining genera were classified as both, either because a single genome contained both specializations or because a genus contained multiple genomes with contrasting specializations. This heterogeneity is best exemplified in *Bilophila* (in the phylum Desulfobacterota) where a majority of the 711 *Bilophila* MAGs were non-TMA producing (97%), 12 MAGs were exclusively proatherogenic, and 7 MAGs encoded both capabilities. While we manually validated this MA content, these analyses were performed on draft genomes of variable completion, thus some MA content could be unsampled. However, since 50% of the dereplicated genomes (n=1,092) in MAGICdb were >90% complete, and classifications were validated for taxonomic consistency, we consider misclassifications due to unsampled genes less likely.

This ability to sample the MA capacity in a genome-wide context highlights a strength of our paired gene and genome databases over prior unpaired single-gene studies. In addition to *Bilophila*, members of the *Desulfovibrio* and a novel genus in Anaerovoracaceae also encoded the proatherogenic choline converting *cutC* along with a TMA reducing *mttB* (**Figure 3D**). This concept of a zero summed game, where there is the potential that TMA is produced and subsequently utilized within the same microorganism, would have been missed where each gene is sampled independently. This finding underscores the value of sampling the entire gene repertoire within a genomic context when identifying microorganisms for possible therapeutic strategies like probiotics.

Since TMA classification largely followed taxonomic lines, it is tempting to want to assign these metabolic roles from taxonomic data alone, as is often done in 16S rRNA amplicon studies in the gut. However, our analyses underlie the danger in doing this, as these metabolisms are not universally encoded by all genome representatives within a genus. For example, of the genomes surveyed, only 51% of the exclusively proatherogenic *Escherichia* and 19% of the non-TMA producing *Blautia* genomes sampled encoded MA metabolism (**Figure 3D**). These analyses reveal the likelihood for falsely reporting an association or metabolic capacity from taxonomic content alone.

In summary, MAGICdb is a high-quality catalog of the TMA modulating genes and genomes that are harbored in the human gut. This curated resource will substantially enhance the sampling precision and efficiency of future microbiome studies. For instance, we show that many of these genomes were rare and not evenly dispersed across humans, thus having a higher likelihood of being missed without cultivation or deep metagenomic sequencing. This extensively sampled reference database can now be used as “bait” to capture this metabolism from less deeply sequenced samples, increasing the ‘mappability’ or recovery of reads for this functional gene content from samples where they would not have assembled. In addition, mapping is a far less computationally intensive process, where users can take advantage of our expertly curated indexing to rapidly annotate this gene content in their datasets.

To demonstrate the useability of MAGICdb in this format, this resource was used in three case studies. We use MAGICdb to map gene expression data from our test fecal reactors and two previously published human cohort studies, illuminating the microorganisms actively shaping TMA concentrations in the gut. We also use this resource to recruit gene content that was previously not sampled in previously published ACVD cohort, demonstrating the efficacy of MAGICdb for disease diagnosing relevance in humans.

### Case study 1: MAGICdb contains microbial members capable of quaternary amine transformations

Previously the active MA transformations in the gut were only demonstrated using pure cultures, thus the cooperative and competitive processing of quaternary amines and their collective contributions to TMA output remains poorly resolved at the community level. To address this knowledge gap, we used fecal reactors to individually dose the same community with each quaternary amine (**Figure 4A**). To uncover the complex metabolic network stimulated by quaternary amines, we profiled the microbial community gene expression with metaproteomics and paired this to quantification of the chemical outcomes with NMR over time.

**Figure 4.**
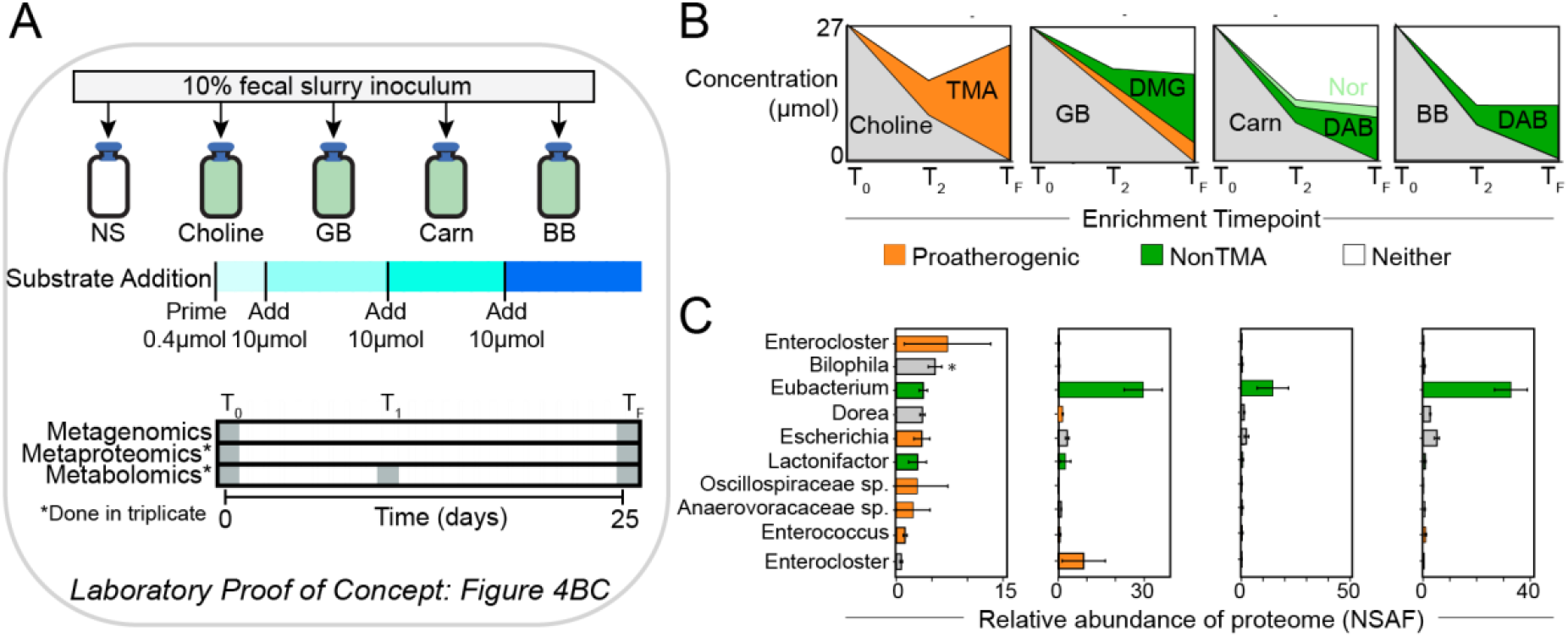
Fecal reactors stimulated with quaternary amines demonstrate MAGICdb contains microorganisms capable of MA transformations. **A** Schematic of fecal reactor study design. Fecal inoculum was provided by an individual in our cohort (see Figure 2B) and stimulated separately with each of the 4 quaternary amines at the dosing shown. Paired multi-omics collected at the beginning and end of the experiment indicated putatively active MA metabolizing microorganisms. **B** Area plots show MA metabolite concentrations in the reactors over time with the curve colored by quaternary amine substrate added (grey) and the microbially produced proatherogenic metabolite TMA (orange) or nonatherogenic metabolite(s) (green) noted. TMA, trimethylamine; DMG, dimethylglycine; Carn, Carnitine; Nor, norcarnitine; DAB, dimethylaminobutyrate; BB, butyrobetaine. **C** The bar chart shows the relative proportion of the metaproteome uniquely assigned to a MAGICdb genome. Microbial bars are colored based on MA gene expression with those potentially contributing to a proatherogenic (orange) or nonTMA producing (green) response denoted. Bars colored in pale gray are genomes that encode MA potential and recruit peptides, but the MA gene content was not expressed under the specific laboratory condition(s). The entire fecal microbial community metaproteome data set is included (**Table S3**).

For both the community wide gene expression and metabolite profiles, replicates within each quaternary amine treatment were congruent, however between treatments the outcomes statistically differed (**Figure S4**). Metabolite quantification revealed that only choline and glycine betaine resulted in a proatherogenic response, while choline was exclusively converted to TMA, glycine betaine conversion resulted in both demethylated and TMA metabolites (**Figure 4B**). Alternatively, carnitine and butyrobetaine stimulation exclusively produced a nonatherogenic response, a finding reflecting the anoxic reactor conditions that restricted the oxygen-requiring proatherogenic monooxygenases (CntA, YeaW)^10,11^.

In the TMA producing reactors amended with choline and glycine betaine, a single genus, *Enterocloster*, dominated the CutC and GrdI expression respectively. Additionally, we provide the first demonstration of TMA production from newly discovered MAGICdb lineages, with genomes from undefined genera in the Anaerovoraceae and Oscillospiraceae implicated in choline conversions. Further the gene expression data supports TMA production from glycine betaine by members of *Dorea, Enterococcus*, and *Enterocloster* genera.

Across the 4 quaternary amine substrates, the nonatherogenic response was mediated by two genomes belonging to the genera *Lactonifactor* and *Eubacterium*, with 3 and 11 *mtxB* genes expressed respectively. This to our knowledge is the first implication for members of the genus *Lactonifactor* in modulating TMA concentrations in the gut. *Eubacterium*, on the other hand, is the model microorganism for quaternary amine demethylation (**Table S1**). Interestingly, we observed that a single *Eubacterium* MtxB was expressed in all quaternary amine treatments, suggesting this single enzyme demethylated all substrates. Validating this supposition, this enzyme was 99% similar to a recently biochemically characterized enzyme purified from *E. limosum* ATCC 8486, which demethylated butyrobetaine, but also promiscuously used carnitine and glycine betaine^15^. Our metabolite and metaproteome findings support a growing body of literature that *Eubacterium* are highly specialized for nonatherogenic quaternary amine degradation in the gut, representing an ideal target for probiotic-based therapeutics.

These genome-based proteome and metabolite analyses also allowed us to contextualize the impacts of MA metabolism more broadly on the gut ecosystem. For instance, the concentration of short chain fatty acids (e.g. acetate, butyrate, propionate) increased across all quaternary amine amended reactors (**Figure S5**). Gene expression data for the genes responsible of the production of these metabolites signaled this was in part due to contributions from MA metabolizing microorganisms (**Figure S6, Table S3**). Given that short chain fatty acids regulate colonocyte energy balance, gut hormone homeostasis, and diabetes^36,37^, gut microbial MA metabolism can have other important health outcomes beyond cardiovascular disease.

This genome context also demonstrated that quaternary amines were metabolized using a variety of energetic strategies. We show that proatherogenic microorganisms process quaternary amines as both a carbon and energy source for anaerobic respiration with fumarate or sulfite as electron acceptors or via obligate fermentation (**Table S3**). The nonTMA producing *Lactonifactor* and *Eubacterium* genomes were inferred to be using these substrates to support and obligatory fermentative lifestyle. Of the ten genomes expressing pro and nonTMA forming genes *Eubacterium* is the only specialist, as all others express glycoside hydrolases with MA genes, such that we cannot rule out concomitant carbohydrate substrate use. Differences in ATP gained from various MA metabolisms will likely impact microbial biomass production and TMA conversion rates, and thus warrant further investigation.

Comparative proteomics indicated that choline and glycine betaine selectively enriched distinct *Enterocloster* strains with non-overlapping substrate specificity (encoded *<u>grdI</u>* or *cutC* but not both) (**Figure 4C**). This exposes the strain-resolved niche differentiation that may occur in the gut. Lastly, even in these reduced complexity, artificially stimulated reactors these data revealed multiple microorganisms co-expressed the same MA enzymes (**Figure 4C**), hinting at the functional redundancy that is likely simultaneously active in the gut.

Together these community focused analyses reveal quaternary amine conversions are an emergent property of the microbiome and cannot be fully evaluated by single gene surveys or outcomes from pure culture experiments. Future engineering of the gut microbiome for controlling TMA concentrations will need to account for energy differences in this metabolism, the metabolic plasticity and exchanges between microorganisms. In summary, moving to an organismal and community resolved view of active MA metabolism sheds light on the complexity that needs to be addressed when designing therapeutic strategies for the gut ecosystem.

### Case study 2: MAGICdb divulges TMA modulating enzymes expressed, but previously obscured in human cohorts

To further extend the relevance of this MAGICdb resource, we mapped human fecal metatranscriptomic and metaproteomic data from two published cohorts containing 306 healthy subjects and 106 subjects diagnosed with irritable bowel disease (**Figure 5A**). We were surprised to find that MAGICdb genes were expressed in 82% of 361 metatranscriptomes and in 58% of 447 metaproteomes (**Figure S7**). Using the metatranscriptome data, we identified the most highly expressed genes for each of the MA gene types and recorded their prevalence (occupancy, % of humans detected in) and mean relative abundance across the cohort (**Figure 5**). We then compared whether these transcribing MA gene containing genera also expressed the same gene in metaproteomes from the human cohort and in our fecal reactors (**Figure 5**).

**Figure 5.**
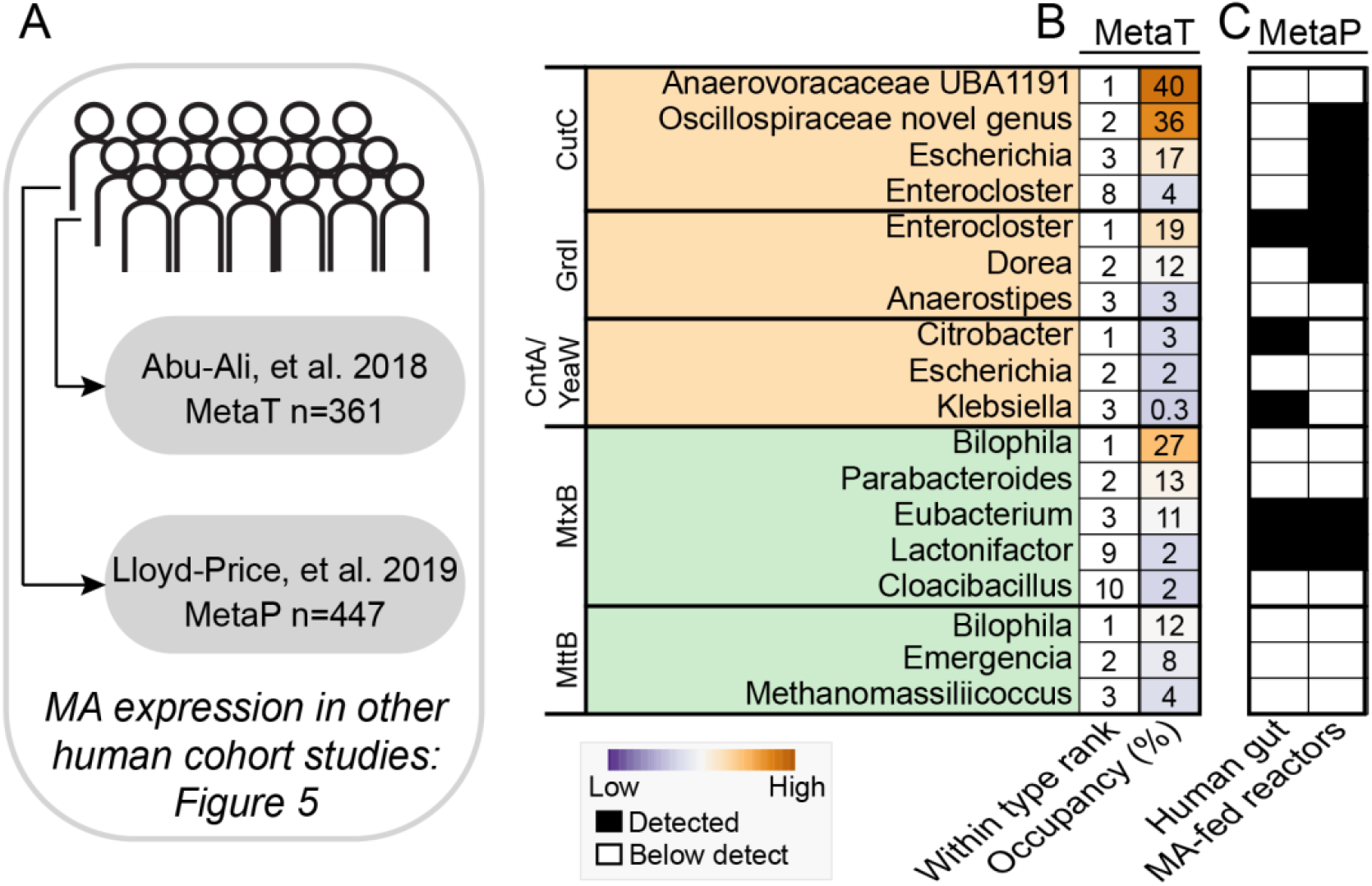
MAGICdb uncovered active microbial members and assigned their metabolic MA roles from *in vivo* human fecal analyses. **A** Schematic showing the use of MAGICdb to recruit expression data from 361 fecal metatranscriptomes (MetaT) collected from a human cohort of 96 individuals and from 447 fecal metaproteomes (MetaP) collected from a human cohort of 75 individuals. **B** For each MA gene type, the top 3 genera with the highest summed gene expression are reported, with some selected lower ranking but genera active in metaproteomes also reported. For each genus the within gene ranking (1 being most expressed) and the cohort occupancy (percentage of metagenomes where gene expression was detected) are quantified. **C** We next compared if these genes were also expressed in metaproteomic data sets derived from our reactors (**Figure 4**) and *in vivo* from a previously published study^64,66^. Shared gene expression data across studies is reported as presence (black) and absence (white).

Across the multitude of data types, fecal sources, and sampled conditions, the same proatherogenic genera were culprits implicated in TMA production from dietary quaternary amines. Our collective findings indicate that the TMA production from choline was likely mediated by microorganisms that have so far evaded cultivation (e.g. members of Anaerovoraceae and Oscillospiraceae). Notably these two genomes had the most highly transcribed *cutC* but were also expressed in more than a third of the 361 metatranscriptome samples (**Figure 5A**). Similarly, we showed members from these genera were active in our choline reactors that exclusively resulted in TMA production (**Figure 4**). While our cohort study revealed that some members of this Anaerovoraceae genus could encode nonatherogenic genes (**Figures 2D, 3B**), our combined *in vivo* and *in vitro* expression data suggest a stronger proatherogenic role may be likely.

Our proatherogenic findings expanded TMA production beyond the Gammaproteobacteria (e.g. *Escherichia, Klebsiella, Citrobacter*) which are well documented to produce TMA, to also include members of the class Clostridia (e.g. *Enterocloster, Dorea*). Notably the exclusively proatherogenic *Dorea* genus (194 genomes in MAGICdb) encoded only *grdI* and the same MAG representative that was active in the human cohort was active in our glycine betaine fecal reactors (**Figure 4C**). Notably, a closely related *Dorea* genome, with the same MA functionality, was the second most dominant in our human cohort and detected in 86% of 52 individuals (**Figure 2D**). While today there is limited progress in designing specific microbiota eradication techniques, our coordinated analyses reveal likely targets for precision interventions.

One of the most significant findings of MAGICdb was our vast expansion of the nonTMA producing microbial enzymes and the microorganisms that encode them. A sequence similarity analyses of the 3,022 MTTB superfamily genes in our database, resulted in 18 clusters composed of 1,031 nodes (**Figure 6**). Forty percent of these sequences were in cluster 1, which contained the genes for directly demethylating proatherogenic TMA. This extends the TMA utilizing gene content (*mttB*) in the human gut from a study focused on a single species of *Bilophila wadsworthia* and a study of TMA utilizing methanogens based on 6 draft genomes^34^. MAGICdb contains 1,071 *mttB* genes assigned to *Bilophila* from multiple species and 61 methanogen genomes that span 3 genera, including one uncultivated genus UBA71 (**Figure 3A**). Outside these lineages, we recovered 407 TMA reducing genomes that were assigned to 7 bacterial genera with considerable representation from members of *Emergencia* (11 MAGs) and a novel genus in the Anaerovoracaceae (2 MAGs) (**Figures 5B, 6**).

**Figure 6.**
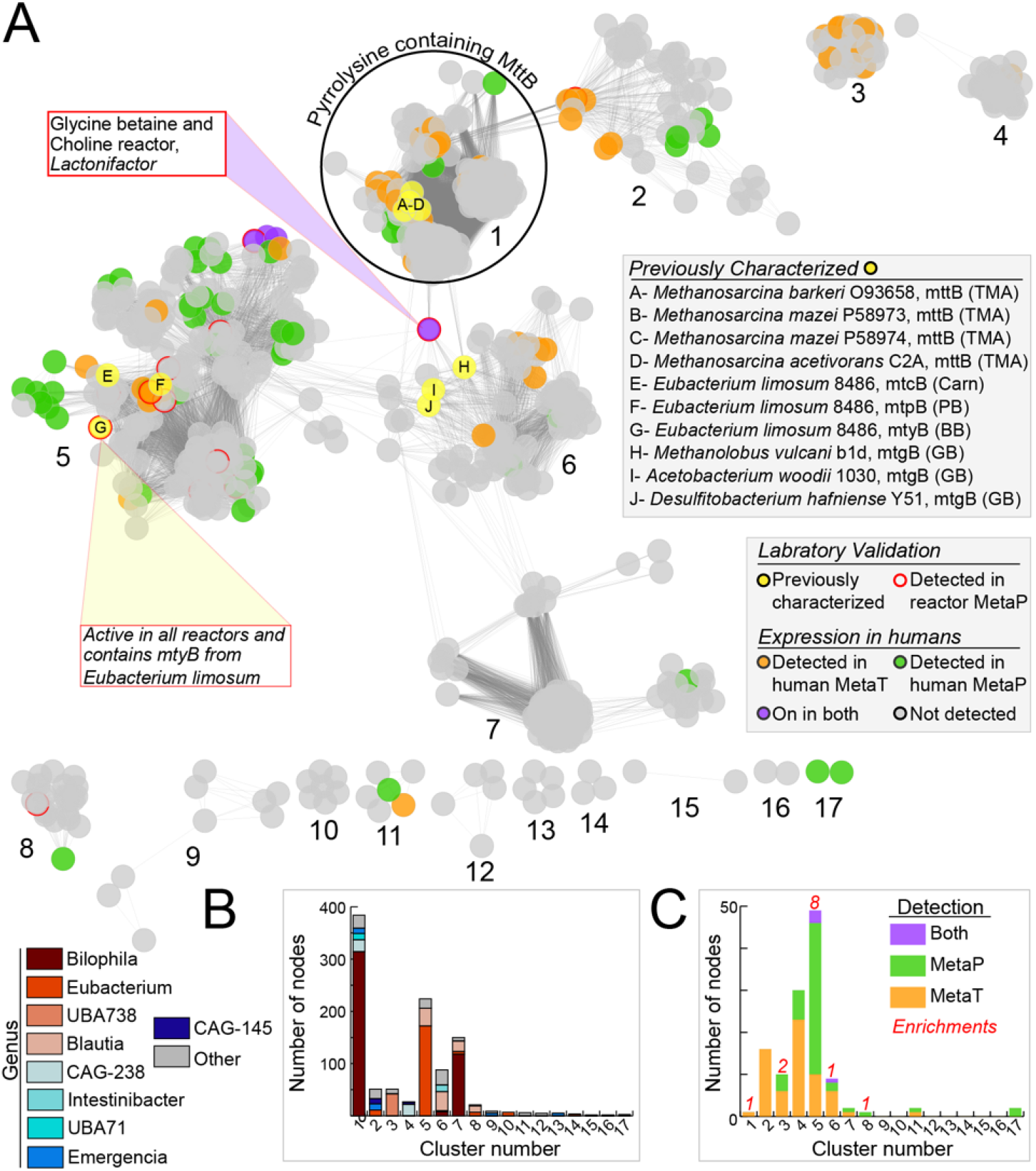
Taxonomy, prevalence, and expression of the MTTB superfamily in MAGICdb. **A** Sequence similarity network (SSN) of the MTTB superfamily within MAGICdb with each of the 1,031 nodes (colored dots) representing one or more amino acid sequences (>99% identity) connected by an edge if the pairwise amino acid sequence similarity is >80%. Nodes are colored to represent gene products that were previously biochemically characterized (yellow filled circles) or recruited to MAGICdb from publicly available microbial gene expression data in feces collected from two large human cohort studies, with metatranscriptome (orange filled circle), metaproteome (green filled circle), or both (purple filled circle) datasets. Nodes with a red outline were expressed in our fecal laboratory metaproteomic data. Previously biochemically characterized MTTB superfamily members are labeled A-J. For these characterized enzymes the microorganisms and preferred substrate are reported in the shaded box with trimethylamine (TMA), Carnitine (Carn), Proline Betaine (PB), Butyrobetaine (BB), and Glycine Betaine (GB) noted. One yellow node, labeled “G”, contained a characterized enzyme from *Eubacterium limosum* ATCC 8486 that was >99% similar to a sequence recovered from a MAG reconstructed here that was expressed in our fecal reactors. Stacked bar charts show the content of each cluster within the SNN, with genus and expression shown in **B** and **C**, respectively.

Here we show members of 7 different genera (e.g., *Bilophila, Emergencia, and Methanomassilicoccus)* expressed 30 of these direct TMA demethylating genes (**Figure 6C**). While an *mttB* gene from a methanogenic *Methanomethylophilaceae* genome was the most highly transcribed (4-fold greater than the others), this gene was only found in 3% of samples. This is true in general for the methanogen TMA demethylating potential, while active in specific humans, it is sparsely distributed (or detected) (**Figure 2E**, *Methanomethylophilaceae* and *Methanomassillicoccus*). The high level of activity of these methanogen and other members in certain humans, which would directly remove TMA from hepatic circulation, indicates how the personalized composition of the gut microbiota between individuals could be an underappreciated moderator of heart disease risk.

This study is the first inventory of nonatherogenic quaternary amine acting gene content (*mtxB*) in the human gut, cataloging 1,863 genes from 43 genera. More than half of the 17 non-pyrrolysine clusters contained a representative that was expressed *in vivo* or *in vitro*, while 2 of these clusters included 6 protein sequences that were previously experimentally verified to demethylate the quaternary amines studied here (**Figure 6**). Of the non-pyrrolysine containing clusters, cluster 5 with 80% of the sequence diversity assigned to *Eubacterium* had the most representatives expressed in human cohorts (**Figure 5**). Interestingly, the *mtxB* gene with the highest mean transcription was assigned to *Bilophila*, which was the genera that most consistently transcribed both *mttB* and *mtxB* across the cohort (**Figure 5**). Beyond known prior roles in TMA production from glycine betaine and choline and direct TMA reduction^34^, we provide evidence that *Bilophila* also has the potential to convert quaternary amines to nonTMA products.

These nonatherogenic routes may control gut TMA concentrations and therefore beneficially contribute to human health. While promising, to determine their translational capacity for probiotic-based therapeutics more research is needed. First, while these genes are inferred to demethylate quaternary amines more in depth biochemical characterization is needed with model organisms beyond *Eubacterium* (**Table S1**). Our combined analyses implicate *Biliophila* or *Parabacteroides* as alternative targets given their activity in human cohorts (**Figure 5**). Also, we showed that many microorganisms containing *mtxB* are not physiologically characterized and would need to be evaluated for their immunogenic potential. Lastly, our genome resolved analyses show the need to audit these demethylating microorganisms for the net TMA production and conversion rate, as these MAGs encode several proatherogenic and nonatherogenic routes. In conclusion MAGICdb recovered MA gene content previously unnoticed in prior microbiome publications, demonstrating the utility of this database to expedite the sampling of microbiome MA metabolism across wider ranges of humans and disease conditions.

### Case study 3: Gut microbiota markers predict cardiovascular disease in humans

While gut microbiota are commonly implicated in cardiovascular disease, the compilation of both proatherogenic and nonatherogenic genes that modulate gut TMA concentrations has not been systematically examined. Previous work sampled the proatherogenic genes that yielded TMA from choline (*cutC*) or carnitine (*cntA*/*yeaW*) in fecal metagenomes from 218 individuals with ACVD and 187 healthy controls. Using a database of 17 genes recovered in the study^32^, this analysis failed to classify disease status in the cohort based on the relative abundance of these genes, with a cross-validation area under the curve (AUC) value of 0.63.

Here we reanalyzed this metagenomic dataset^32^, but instead used the 3,031 unique genes in the MAGIC database for read recruitment (**Figure 3A**). For context, MAGICdb has a 62- and 161-fold more sampling of *cutC* and *cntA*/*yeaW* gene richness respectively, but also included the other MA genes not in the previous analysis (*grdI, mtxB, mttB*). Through read mapping MAGICdb uncovered 2,699 unique MA genes residing in these fecal metagenomes (**Figure S7)**. We showed that ACVD subjects had increased relative abundance of proatherogenic genes (*cutC, cntA, yeaW, grdI*), while the nonatherogenic *mttB* gene relative abundance was depleted in these same individuals (**Figure 7A**).

**Figure 7.**
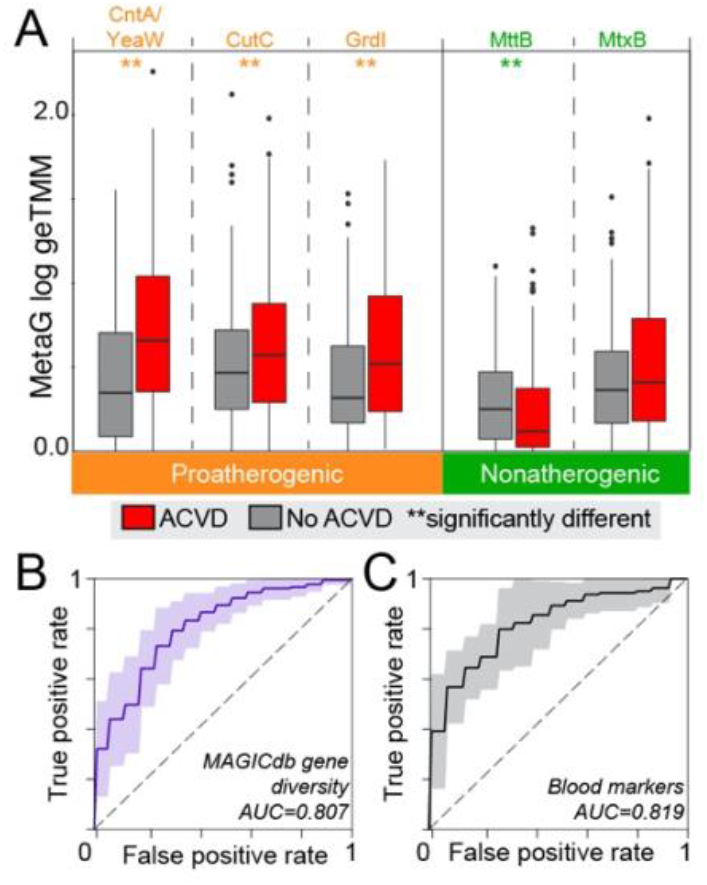
MAGICdb gene content predicted cardiovascular disease in humans. **A** Boxplots display the relative abundance of MAGICdb genes in fecal metagenomes from ACVD patients (red) or non-ACVD control subjects (grey), with significant differences by ACVD status denoted by double asterisk. Receiver operating curves show the area under curve (AUC) for predictions of ACVD status in humans by **B** richness and distribution of MAGICdb genes and **C** blood markers (LDL, HDL, and triglycerides). These microbiome and host derived AUC were not significantly different (McNemar’s, p-value >0.05).

To ascertain the enhanced prediction provided by the increased gene richness, the logistic regression model using only *cutC* and *cntA/yeaW* from MAGICdb had an AUC value of 0.67, with a slightly improved classification from the original model (**Figure S8**). However, expanding our model to include the relative abundance for the full gene set (*cutC, cntA, yeaW, grdI, mtxB*, and *mttB*) and the diversity profile of these genes across the cohort (**Figure 7B**) added to the predictability, resulting in AUC values of 0.75 and 0.81 respectively. These significantly increased values indicated the power of including both proatherogenic and nonatherogenic gene diversity when classifying ACVD health status. With an AUC of 0.81 these MA gene-based predictions did not differ significantly from predictions reliant on more traditional cumulative blood markers (HDL, LDL, triglycerides) in this same cohort (**Figure 7C**). Future studies with larger cross-sectional cohorts of ACVD and healthy individuals are required to validate this microbiome derived gene model more completely.

### Conclusion

Tradeoffs between the individuals sampled and sequencing depth impact all cohort studies, and are important considerations when functional genes, especially those encoded by rare members need to be sampled. Our hope in creating MAGICdb was to alleviate this burdened decision from future cardiovascular disease relevant cohort studies. We have demonstrated that expertly curated MAGICdb eliminates the need to individually deeply sequence to assemble and recover these rare, sparsely distributed genes, instead favoring shallower sequencing from more individuals, with MA genes recovered by mapping to the database.

MAGICdb is the first comprehensive catalog of gut microbial MA metabolism. We leveraged this database with our own and other’s metagenome, metatranscriptome, and metaproteome datasets to show that these gut MA genes are active, coordinated, and predictive of cardiovascular disease. This open-access database and accompanying models can be applied to larger cohorts, opening the door for nontraditional microbiota tools for diagnosing, halting, and reversing cardiovascular disease. Additionally, this gene foundation can be exploited to discover the microbial MA contributions to other diseases (e.g diabetes, as well as cerebral, hepatic, and vascular conditions) where this metabolism has been implicated^38,39^. Collectively, our results establish the utility of metabolism-oriented microbiome databases to guide modern precision medicine strategies designed to correct defects in the gut microbiome.

## Materials and Methods

### MAGICdb construction and analysis

Combining the 1,436 MAGs recovered in this study with (i) 700 genomes from isolates in the Human Microbiome Project (HMP)^26^ and (ii) 237,273 gut derived metagenome-assembled genomes (MAGs) from previously published studies^24,25^, we obtained 238,530 gut associated genomes for analysis of MA metabolic potential. MAGs in (ii) were compilation studies, where MAGs were accumulated across many publications representing many different lifestyles, disease types, and diets^24,25^. As outlined in **Figure S1**, each gene type in **Figure 1** was assessed separately. First, using an experimentally validated amino acid sequence, each gene type was searched against the predicted amino acid sequences of the 238,530 gut associated genomes using BLAST^40^, retaining sequences with >60 bitscore. For CutC, CntA, YeaW, and GrdI, sequences were aligned with experimentally validated reference sequences using muscle, and phylogenetic trees were built using RAxML^41^. Individual gene trees were visualized in iTOL^42^, and the branch containing sequences of interest were selected. For the remaining sequences, active residues were confirmed as outlined for CutC, CntA, YeaW, and GrdI^8,10,11,21^. Of note, is CntA and YeaW, which we report together as specificity cannot be inferred from sequence information alone ^10,11^. The remaining sequences with active residues were then incorporated into MAGIC gene database, as well as their corresponding genomes into MAGIC genome database.

For MttB superfamily genes that did or did not contain pyrrolysine, a different approach was taken due to pyrrolysine interpreted as a stop codon during gene calling^18,19^. After recovery of putative MttB homologs using amino acid BLAST^40^, obtained sequences were length filtered to 360 bp and aligned to known MttB superfamily members. Sequences longer than 360 did not contain pyrrolysine and aligned through the pyrrolysine residue were incorporated into the MAGIC gene database as non-pyrrolysine containing MtxB, as well as their corresponding genomes into the MAGIC genome database. These superfamily sequences could not be assigned a specific quaternary amine substrate, as such we denoted these as MtxB to indicate an unassigned substrate “X”, nomenclature consistent with the MttB superfamily (e.g. MtgB for glycine betaine^6^, MtcB for carnitine^13^). The remaining truncated genes were then manually called in Geneious^43^ from the original genome scaffolds using the amber read-through option to detect pyrrolysine. The resulting sequences that encoded for pyrrolysine were incorporated into the MAGIC gene database as pyrrolysine containing MttB, as well as their corresponding genomes into the MAGIC genome database.

MTTB superfamily genes in MAGICdb were used to construct a sequence similarity network via the EFI-EST webtool^44^. Networks were generated with initial edge values of >80%, and sequences with 100% sequence similarity were collapsed into single nodes. The resulting representative node network was visualized with Cytoscape 3.8^45^ using the perfuse force directed layout option. Genomes in MAGICdb were analyzed with GTDB-Tk^46^ for taxonomy, checkM^47^ for quality, and DRAM^48^ for genome annotation.

### Sample procurement and cohort statistics

The current study considered samples collected from 125 individuals aged 21 years or older under the auspices of Dr. Alan George Smulian either at the University of Cincinnati College of Medicine or the University of Cincinnati Medical Center Holmes Hospital Outpatient Services. Each individual provided self-collected fecal and urine samples, along with data on medical history (e.g. antibiotic usage, recent colonoscopy), weight, age, dietary habits, and smoking status (**Table S1**). Donor identities were stripped from the paired samples and their associated data, and each donor was assigned a unique identification number. Targeted metabolomic analyses of methylated amines (MAs) were carried out on fecal samples from all 125 individuals, while a subset of 54 samples were selected based for fecal metagenomic sequencing along a fecal TMA gradient (**Figure S3**). Based on surveys, subjects and their corresponding samples were removed from analyses due to antibiotic use in the last 6 months, lack of patient information, or a colonoscopy in the last 6 months, confining the cohort to 113 subjects. Five sets of donated samples were removed from analyses due to donor antibiotic use and seven were removed for lack of donor de-identified data. Written, informed consent was obtained from all study participants, and subject treatment and experiments with donated samples were approved by Institutional Review Boards of the University of Cincinnati and the Ohio State University.

### Metagenomic sequencing, assembly, and binning for this cohort and methylated amine reactors

Fifty-four fecal samples out of 113 were chosen across a trimethylamine gradient for metagenomic sequencing, with at least five samples chose from each quartile (**Figure 2B**). Total nucleic acids were extracted from five microcosm samples and 54 human fecal samples using the PowerSoil DNA Isolation kit (MoBio), eluted in 100 μL, and stored at −20 °C until sequencing. DNA was submitted for sequencing at the Genomics Shared Resource facility at The Ohio State University. Libraries were prepared with the Nextera XT Library System in accordance with the manufacturer’s instructions.

Genomic DNA was sheared by sonication, and fragments were end-repaired. Sequencing adapters were ligated, and library fragments were amplified with five cycles of PCR before solid-phase reversible immobilization size selection, library quantification, and validation. Libraries were sequenced on the Illumina HiSeq 2500 platform and paired-end reads of 113 cycles were collected. All raw reads from microcosms and fecal samples were trimmed from both the 5′ and 3′ ends with Sickle (https://github.com/najoshi/sickle), and then each sample was assembled individually with IDBA-UD^49^ using default parameters. Metagenome statistics including amount of sequencing are noted in **Table S1**.

All microcosm and fecal metagenomes (**Table S1**) were binned using metabat2^50^ with default parameters. Bins were then assessed for quality using checkM^47^. Metagenomic reads from the binned samples were then mapped to bins >50% completion and <10% contamination (medium or high quality bins^51^) at 99% identity using bbmap^52^. For deeply sequenced metagenomes (n=15) reads that did not map to the pool of medium or high quality bins were then reassembled using IDBA-UD^49^, completing iterative assemblies for each of the 15 samples, until no new bins could be recovered. The resulting 2,447 bins were then dereplicated into 1,436 bins using dRep (*55*).

### Fecal metabolite analyses from the cohort study

Fecal samples were self-collected by volunteers and brought to the collection center where they were stored at −80 °C. Samples were then shipped to the lab for analysis on dry ice where they were again stored. Samples arrived frozen in less than 24 h and were immediately stored at −80 °C until ready for NMR analysis.

Fecal samples were removed from the freezer and transferred to a biosafety cabinet on dry ice. A total of 0.2 to 0.5 g (wet weight) of frozen chips of each sample were weighed and transferred to a 5 ml centrifuge tube. To extract metabolites from the fecal samples, 1 ml 0.75 M potassium phosphate buffer (PBS buffer) in 50% D2O, pH 7.2, was added to each tube, resulting either 3x volume/weight dilution (for fecal samples with more than 0.3 g in wet weight) or 5x volume/weight dilution (for fecal samples with less than 0.3 g in wet weight) of the original samples. The slurries were then vortexed for a total of 3 minutes to extract metabolites. Vortexing was paused several times in order to cool the sample on ice to avoid overheating. The vortexed samples were then centrifuged at 1000x g for 10 minutes at 4°C. The supernatant was transferred to a 1.5 ml microcentrifuge tube and were centrifuged again twice at 4°C (16100x g, 10 min) to remove remaining debris. Total 200 ul of final supernatant were mixed with 100 uM DSS and transferred to a 3 mm x 178 mm NMR tube for NMR analysis.

1D ^1^H and 2D ^1^H-^13^C HSQC NMR spectra were conducted at 298 K on a Bruker Avance III HD 800 MHz (Billerica, MA) at Ohio State campus chemical instrument center (CCIC) NMR facility. Proton NMR, about 4 min for one Data Set, was acquired using 1.28s acquisition time, 2s relaxation delay, and 64 number of scans. The water suppression was achieved using excitation sculpting with gradients. 2D ^1^H-^13^C HSQC was acquired with a standard Bruker pulse sequence using phase-sensitive echo/antiecho-TPPI gradient selection. The experiment parameters include ∼4ms acquisition time in ^13^C dimension, ∼80ms acquisition time in ^1^H dimension, 1s relaxation delay, 16 number of scans, ^13^C GARP decoupling during acquisition, and data matrix of 2048 × 128. The experimental time is roughly 38 min for one Data Set. Standards with 100 uM of target metabolites (>98% purify) were analyzed under the same conditions. When appropriate, sample aliquots were spiked with a known concentration of trimethylamine to confirm peak assignment.

All NMR data were processed with Bruker Topspin 3.6.1 (Billerica, MA). The data were typically zero-filled one time in both ^1^H and ^13^C dimension prior the application of window functions, followed by Fourier transformation, phasing, and baseline correction. Chemical shifts were internally referenced to DSS at 0.00 ppm. The concentration of a trimethylamine was estimated employing standards of known concentration and comparing the integral of peaks to DSS.

### Cohort analyses

We leveraged our cohort metagenomes to understand the distribution of MA genes with variable depths of sequencing and in relation to fecal TMA concentrations. First, we mined our fecal metagenome assemblies for MA genes, finding 153 MA genes that were dereplicated into 135 genes using cd-hit^53^. We grouped subjects into quartiles (Q1-Q4, 25% of the data points in each) and then related the paired gene content as shown in **Figure 2B**. To understand the recovery of new genes with additional subjects and sequencing, we performed a species accumulation analysis where genes recovered from each metagenome were iteratively dereplicated with the addition of each subject using cd-hit^53^, as shown in **Figure 2C**. To obtain gene abundance, we mapped metagenomic reads rarified to 8Gbp of sequencing to the dereplicated gene set (n=135) using bowtie2^54^. Reads were counted and summarized using coverM (https://github.com/wwood/CoverM) into trimmed mean (-m trimmed_mean) and including genes with a minimum covered fraction of 75% (--min-covered-fraction 0.75). To relate gene abundance to fecal TMA concentration, we used linear regression based modelling to predict TMA concentrations from MA gene relative abundance in our cohort using sparse Partial Least Squares (sPLS ^55,56^) as implemented in the R package mixOmics^57^, with data shown **in Figure S2D**. To further understand how gene recovery was impacted by sequencing depth, we used our most deeply sequenced metagenomes (>35Gbp) to recruit reads to the cohort gene database (n=135) using all reads for a particular metagenome and reads rarified to 4Gbp from the same metagenome, with gene abundance and gene count reported in **Figure S2C**. Briefly, we mapped all metagenomic reads or reads rarified to 4Gbp (similar to previously depths used in other microbiome studies^27,32^) of sequencing to the dereplicated gene set (n=135) using bowtie2^54^. Reads were counted and summarized using coverM (https://github.com/wwood/CoverM) into trimmed mean (-m trimmed_mean) and including genes with a minimum covered fraction of 75% (--min-covered-fraction 0.75).

Beyond the gene level, within our cohort, we aimed to understand the distribution of MA genomes in the context of the microbial community. Abundance data reported was based on the 1,436 unique bins. Briefly, reads from metagenomes with greater than 8Gbp in depth were rarified to 8Gbp from all 52 metagenomes and mapped to 1,436 unique bins using bowtie2^54^ with 95% identity and counted using coverM (https://github.com/wwood/CoverM) with in trimmed mean mode (-m trimmed_mean) and including genomes with a minimum covered fraction of 75% (--min-covered-fraction 0.75). Trimmed mean values were then transformed into relative abundance. To determine rank of each genome, relative abundance of each genome was averaged across the cohort and then ordered from maximum to min, and ranks were assigned 1-1,436, as shown in **Figure 2DE**. Note, only 52 metagenomes were used in this analysis, as two were dropped due to sequencing <8Gbp.

### Methylated amine reactor construction and operation

The microcosm experiment consisted of six treatments all set up with fecal material from subject 74: (i) no substrate and fecal material, (ii) glycine betaine and fecal material, (iii) carnitine and fecal material, (iv) butyrobetaine and fecal material, and (v) choline and fecal material. Each treatment was done in triplicate and consisted of 10% (wet weight/volume) anoxic, fecal slurry in sterile basal bicarbonate-buffered medium dispensed in Balch tubes sealed with butyl rubber stoppers and aluminum crimps under an atmosphere of N_2_/CO_2_ [80:20 (vol/vol)], with a final volume of 10mL. Before mixing with fecal slurry, the medium (per liter) included 0.25 g ammonium chloride, 0.60 g sodium phosphate, 0.10 g potassium chloride, 2.5 g sodium bicarbonate, 10 ml DL-vitamin mixture, and 10 ml DL-mineral mixture and was brought to a pH of 7.0 using 1 mM NaOH^58^. Tubes were incubated at 37°C. Samples for metagenomics and metaproteomics were taken at the final (T_F_) timepoint, while metabolite samples were taken at the indicated times during the course of the 25-day incubation (**Figure 4A**). Anoxic fecal reactors were primed with 40μM of each substrate from time of inoculation to day 3, then they were dosed with 1mM of each substrate 3 times at day 3, day 10, and day 17. Accounting for removal of 1mL samplings, a total of 27umol of each substrate was added. Samples were taken for subsequent analysis at T1 (10 days), T2 (17 days), and TF (25 days). For timepoints T1 and T2, samples were taken prior to substrate addition. Subject 74 fecal material, used for reactor inoculum, metabolite concentrations are given in **Table S3**.

### Methylated amine reactor metabolomic data acquisition and analysis

Samples from microcosm experiments were filtered (0.2 μm) at time of collection and sent to the Pacific Northwest National Laboratory for metabolite analysis by NMR. Samples were diluted by 10% (vol/vol) with 5 mM 2,2-dimethyl-2-silapentane-5-sulfonate-d_6_ as an internal standard. All NMR spectra were collected using a Varian Direct Drive 600-MHz NMR spectrometer equipped with a 5-mm triple resonance salt-tolerant cold probe. The 1D ^1^H NMR spectra of all samples were processed, assigned, and analyzed using Chenomx NMR Suite 8.3 with quantification based on spectral intensities relative to the internal standard. Candidate metabolites present in each of the complex mixtures were determined by matching the chemical shift, J-coupling, and intensity information of experimental NMR signals against the NMR signals of standard metabolites in the Chenomx library. The 1D ^1^H spectra were collected following Chenomx data collection guidelines^59^, using a 1D NOESY presaturation (TNNOESY) experiment with at least 512 scans at 298K using a 100ms mixing time, with 12 ppm spectral width, a 4s acquisition time followed by a relaxation delay of 1.5 s during which a presaturation of the water signal applied. Post-acquisition processing included time domain free induction decays (57472 total points) zero-filling to 132k points and multiplication by a decaying exponential function (line broadening of 0.5 Hz) prior to Fourier Transform. Chemical shifts were referenced to the ^1^H methyl signal in DSS-d_6_ at 0 ppm. Additionally, 2D spectra (including ^1^H–^13^C heteronuclear single-quantum correlation spectroscopy, ^1^H-^1^H total correlation spectroscopy) were acquired on a subset of the fluid samples. Biological triplicates had similar metabolite pools, with all data reported (**Table S3**).

### Methylated amine reactor metaproteomic extraction, spectral analysis, and data acquisition

Liquid culture (1.2 mL) from each microcosm sample was collected anaerobically, centrifuged for 15 min at 10,000 × g, separated from the supernatant, and stored at −80 °C until shipment to Pacific Northwest National Laboratory. Proteins in the pellet were precipitated and washed twice with acetone. Then the pellet was lightly dried under nitrogen.

Each precipitated protein pellet was diluted in 200 μL of 8 M urea in 100 mM ammonium bicarbonate, pH 8 (ABC). A bicinchoninic acid (BCA) assay (Thermo Scientific, Waltham, MA USA) was performed to determine protein concentration. Following the assay, 10 mM dithiothreitol (DTT) was added to the samples and incubated at 60°C for 30 mins with constant shaking at 800 rpm. Samples were then diluted 8-fold for preparation for digestion with 100 mM ABC, 1 mM CaCl2 and sequencing-grade modified porcine trypsin (Promega, Madison, WI) was added to all protein samples at a 1:50 (w/w) trypsin-to-protein ratio for 3 h at 37°C. Digested samples were desalted using a 4-probe positive pressure Gilson GX-274 ASPEC™ system (Gilson Inc., Middleton, WI) with Discovery C18 50 mg/1 mL solid phase extraction tubes (Supelco, St.Louis, MO), using the following protocol: 3 mL of methanol was added for conditioning followed by 2 mL of 0.1% TFA in H2O. The samples were then loaded onto each column followed by 4mL of 95:5: H2O:ACN, 0.1% TFA. Samples were eluted with 1mL 80:20 ACN:H2O, 0.1% TFA. The samples were concentrated down to ∼100 μL using a Speed Vac and a final BCA was performed to determine the peptide concentration and samples were diluted to 0.1 μg/μL with nanopure water for MS analysis.

All mass-spectrometric data were acquired using a Q-Exactive Plus (Thermo Scientific) connected to a nanoACQUITY UPLC M-Class liquid chromatography system (Waters) via in-house 70-cm column packed using Phenomenex Jupiter 3-μm C18 particles and in-house built electrospray apparatus. MS/MS spectra were compared with the predicted protein collections using the search tool MSGF+^60^. Contaminant proteins typically observed in proteomics experiments were also included in the protein collections searched. The searches were performed using ±20-ppm parent mass tolerance, parent signal isotope correction, partially tryptic enzymatic cleavage rules, and variable oxidation of methionine. In addition, a decoy sequence approach^61^ was employed to assess false-discovery rates. Data were collated using an in-house program, imported into a SQL server database, filtered to ∼1% false-discovery rate (peptide to spectrum level), and combined at the protein level to provide unique peptide count (per protein) and observation count (that is, spectral count) data. Spectral count data for each identified protein was normalized using normalized spectral abundance frequency (NSAF) calculations^62,63^, accounting for protein length and proteins per sample (**Table S3**). Note that metaproteomics were not done on raw fecal samples. Metaproteomes were mapped to dereplicated MAGICdb predicted amino acid sequences, as well as predicted amino acid sequences of unique MAGs recovered from enrichments.

### Mapping of published data to MAGICdb

All reads were downloaded from EBI from Abu-Ali, et al.^64^, a study of metatranscriptomes from adult men. Adapters were stripped using bbduk.sh with the parameters ktrim=r, k=23, mink=11, hdist=1. Reads were trimmed using sickle with default parameters. Reads were mapped to MAGICdb genes using bbmap.sh (bbtools suite^52^) using perfectmode=t and ambiguous=random. Counts were extracted from the bbmap covstats output and compiled into a table. The counts were then transformed to geTMMs^65^.

All proteome .mgf files were downloaded from Lloyd-Price, et al.^66^. Files were then searched against the MAGICdb using MSGF+^60^ using the parameters inst 3, tda 1, ti 1,3, ntt 1 and maxLength 50. After the search files were converted to TSVs using the parameter showDecoy 1. To determine hits, first all hits with a pep q-value greater than .01 were removed. Then for each sample we identified proteins with more than one peptide hit. This list of proteins per sample were the ones considered present.

### CVD prediction from human gut metagenomic data

All reads were downloaded from EBI from Jie, et al.^32^, a study of metagenomes from 218 individuals with atherosclerotic cardiovascular disease (ACVD) and 187 healthy controls (*15*). Adapters were stripped using bbduk.sh with the parameters ktrim=r, k=23, mink=11, hdist=1. Reads were trimmed using sickle with default parameters. Reads were mapped to unique MA genes in MAGICdb genes using bbmap.sh (bbtools suite^52^) using perfectmode=t and ambiguous=random. Counts were extracted from the bbmap covstats output and compiled into a table. The counts were then normalized to geTMM, a gene length corrected (ge) trend means of m-values (TMM), which is a method for assessing intrasample variation for read map data^65^. For each individual we obtained the relative abundance profile of the MA genes (*cutC, cntA, yeaW, grdI, mtxB*, and *mttB)*.

The relative abundance MA gene profiles were then used in a logistic regression model using scikit-learn^67^ to predict ACVD status (0=No ACVD, 1=ACVD) as designated in Jie, et al^32^. Models were evaluated using stratified 10-fold cross-validation with mean false positive and true positive rates reported and used to calculate the area under the receiver operator characteristic curve (AUC-ROC)^68^. Feature coefficients for logistic regression model for the best performing model (Shannon’s diversity of each type of gene in MAGICdb) were reported. Gender was included in the models were noted and dichotomized so that male equals 1 and female equals 0. An AUC value >0.7 was used to indicate a relatively good ability for the model to classify individual disease status^69^. To test for difference in model performance McNemar’s test was used^70^.

Models were trained on the following to predict ACVD status:

1. Shannon’s diversity of MAGICdb genes by gene type + gender (**Fig 7B**). Shannon diversity score was determined for each gene type using the scikit-bio (*http://scikit-bio.org/*) and calculated using the geTMMs. Each individual had a Shannon’s diversity profile that included 6 MA gene diversity scores per individual.
2. Blood markers (triglyceride mmol/L, LDL mmol/L and HDL mmol/L) + gender (**Fig 7C)**
3. Abundance of *cutC, cntA/yeaW* summed per gene type + gender (genes used and model analysis similar to reported in Jie, et al^32^ (**Fig S8A**)
4. Abundance of all genes from MAGICdb + gender (**Fig S8B**), each gene abundance in the unique MAGICdb gene database is included.
5. Abundance summed per gene type + gender (**Fig S8C**)
6. Abundance of all genes summed per atherogenic status (proatherogenic and nonatherogenic) + gender (**Fig S8D**).
7. Shannon’s diversity of MAGICdb genes by gene type (no gender) (**Fig S8E**). Similar to above in number 1 but did not include gender.

## Data availability

All data will be made available upon submission.

---

**Figure S1.**
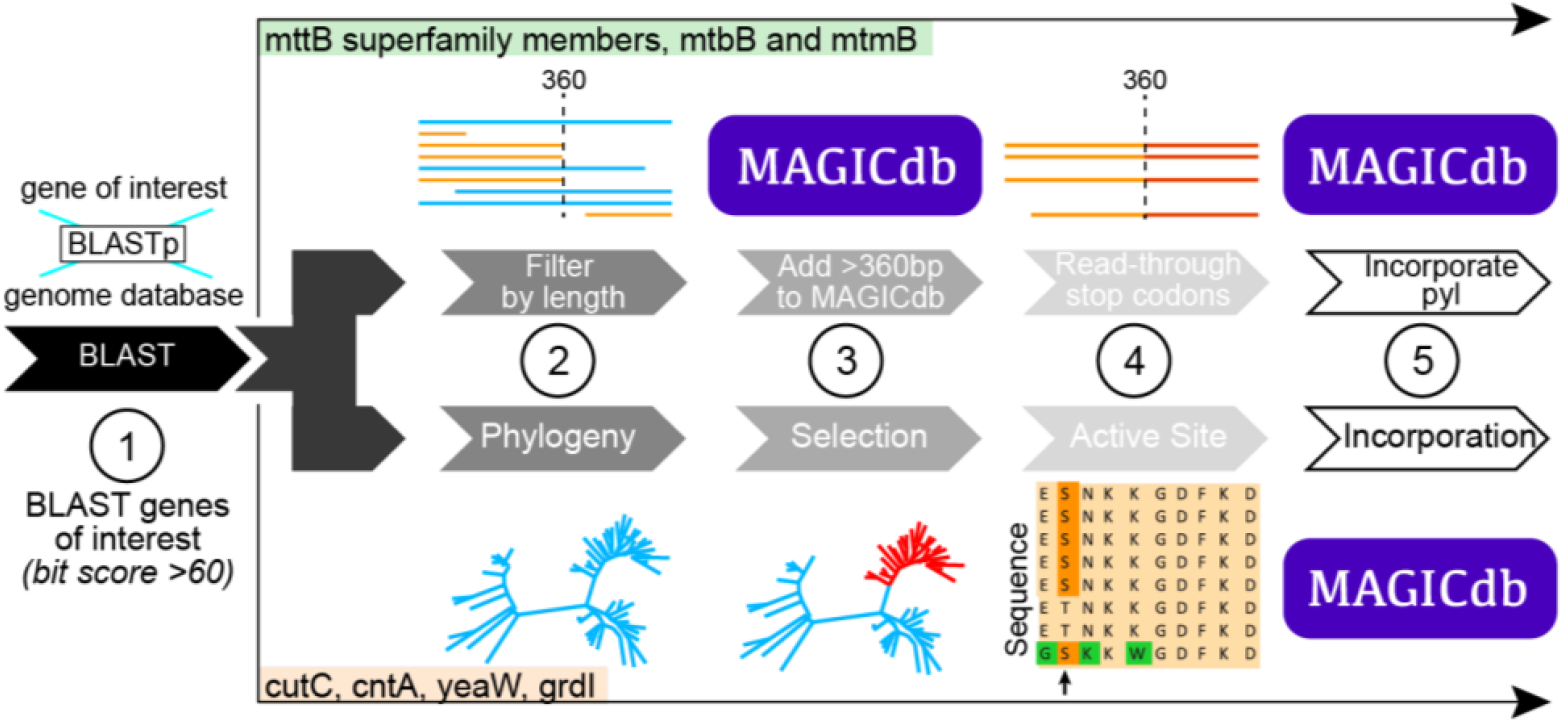
The computational workflow for MAGICdb uses both homology and nonhomology based approaches to refine the annotation of MA gene content. Details can also be found in methods section.

**Figure S2.**
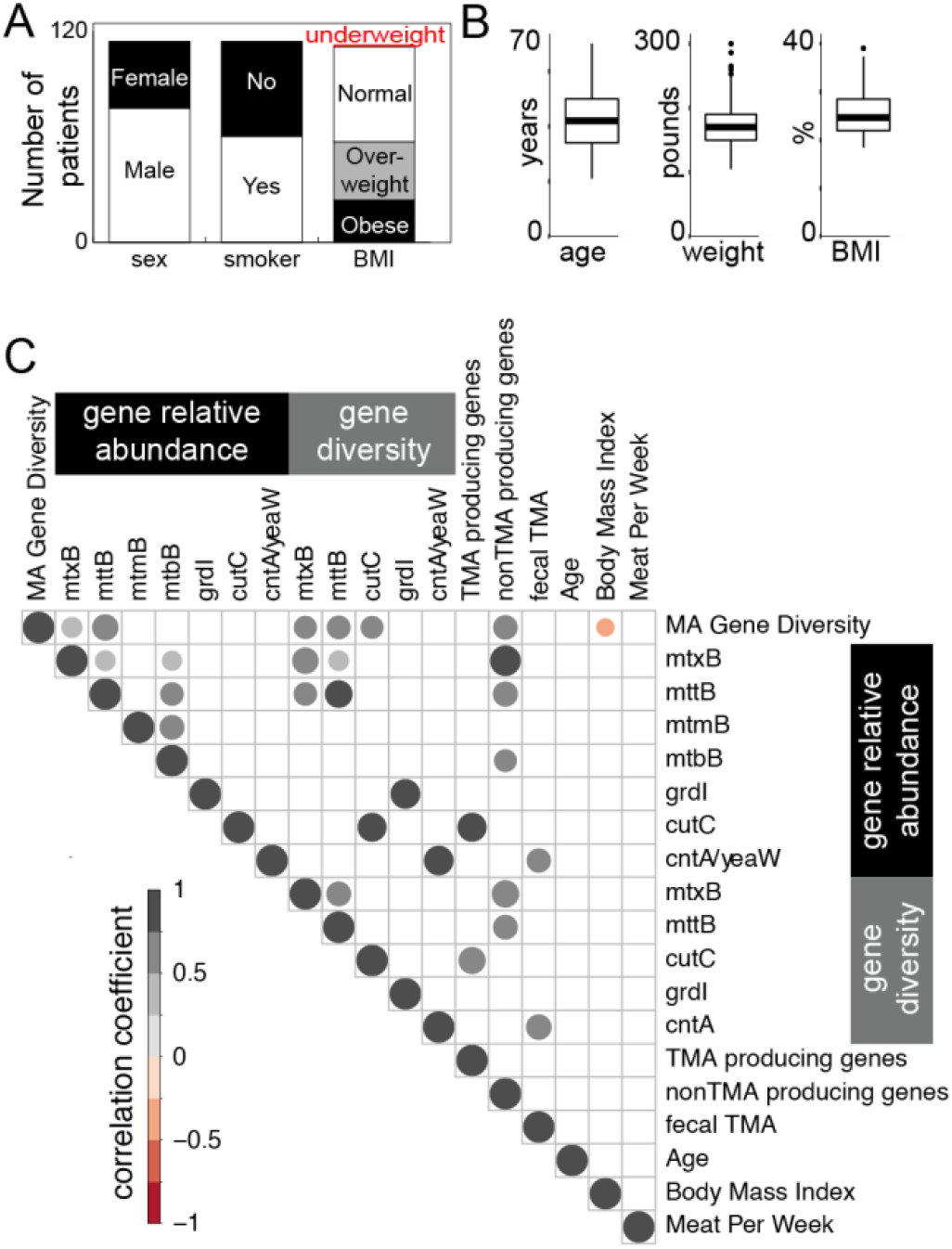
**A** Cohort statistics including sex, smoking status, and BMI category of 113 human subjects. **B** The median values of age, weight, and BMI statistics across the cohort (n=113). Points above or below boxplots signify outliers, or values outside one standard deviation of the median. **C** Dot plot shows the all-to-all correlations of MA gene diversity, MA gene abundance, and host lifestyle factors, with the significant (p-value<0.01) correlations shown by dots colored and sized by correlation coefficients.

**Figure S3.**
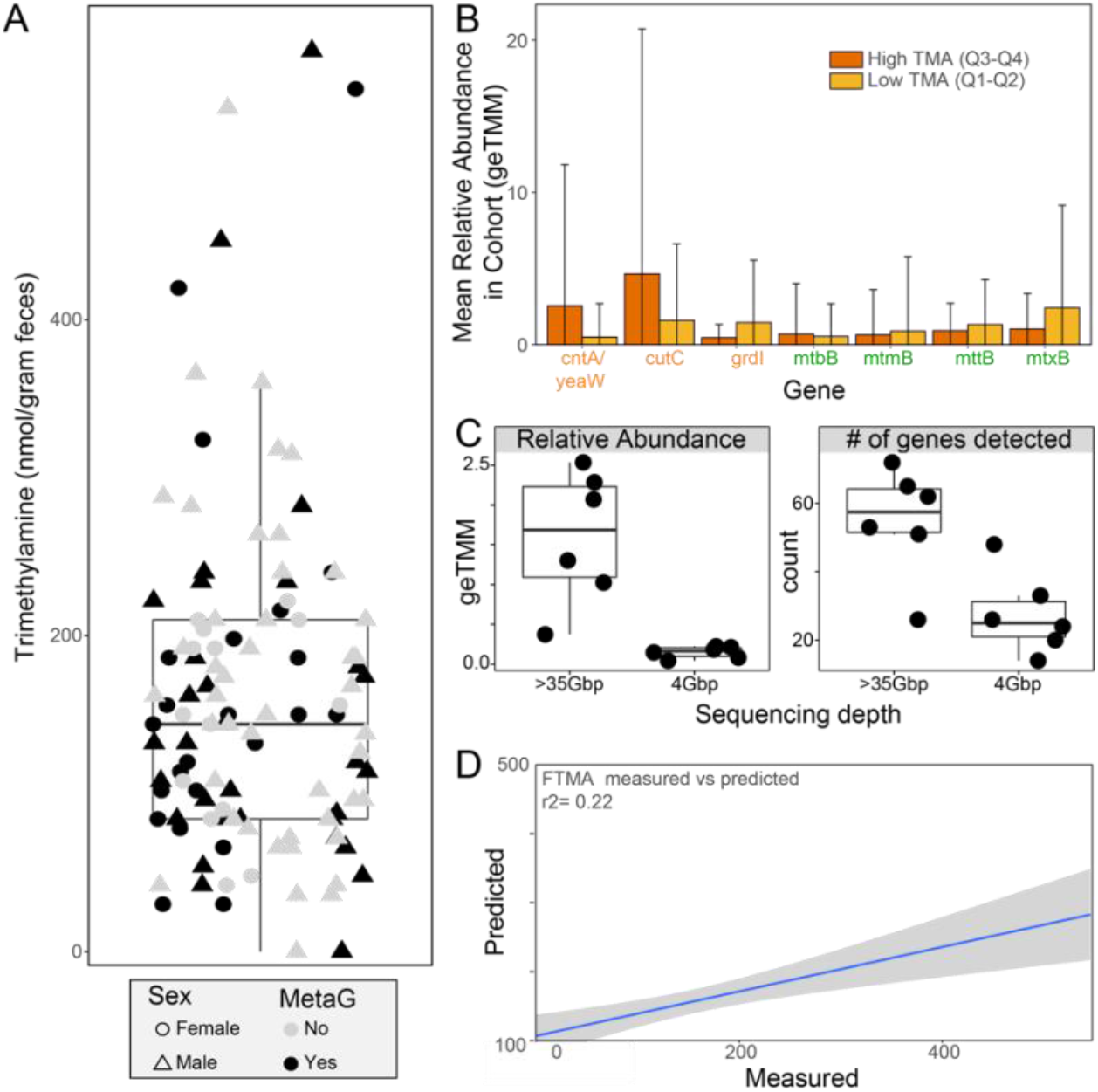
**A** Fecal TMA concentrations, with color denoting the samples sequenced with metagenomics (n=54, black) and those not chosen (gray). Shape indicates the sex of the subject from which the fecal sample was derived. **B** Rarified metagenomes (8Gbp) were mapped to a database of 135 dereplicated MA genes to provide the mean relative abundance of each gene type across the cohort. The mean and standard deviation of the relative abundance of MA genes recovered from high (>145 nmol TMA/gram feces) and low (0-144 nmol TMA/gram feces) fecal TMA samples. **C** Comparison of the relative abundance (right) and count of MA genes recovered when >35Gbp and 4Gbp of reads were evaluated. **D** Prediction of fecal TMA from MA gene content using sparse Partial Least Squares (sPLS) regression revealed a significant relationship between the MA gene content predicted and measured fecal TMA concentrations.

**Figure S4.**
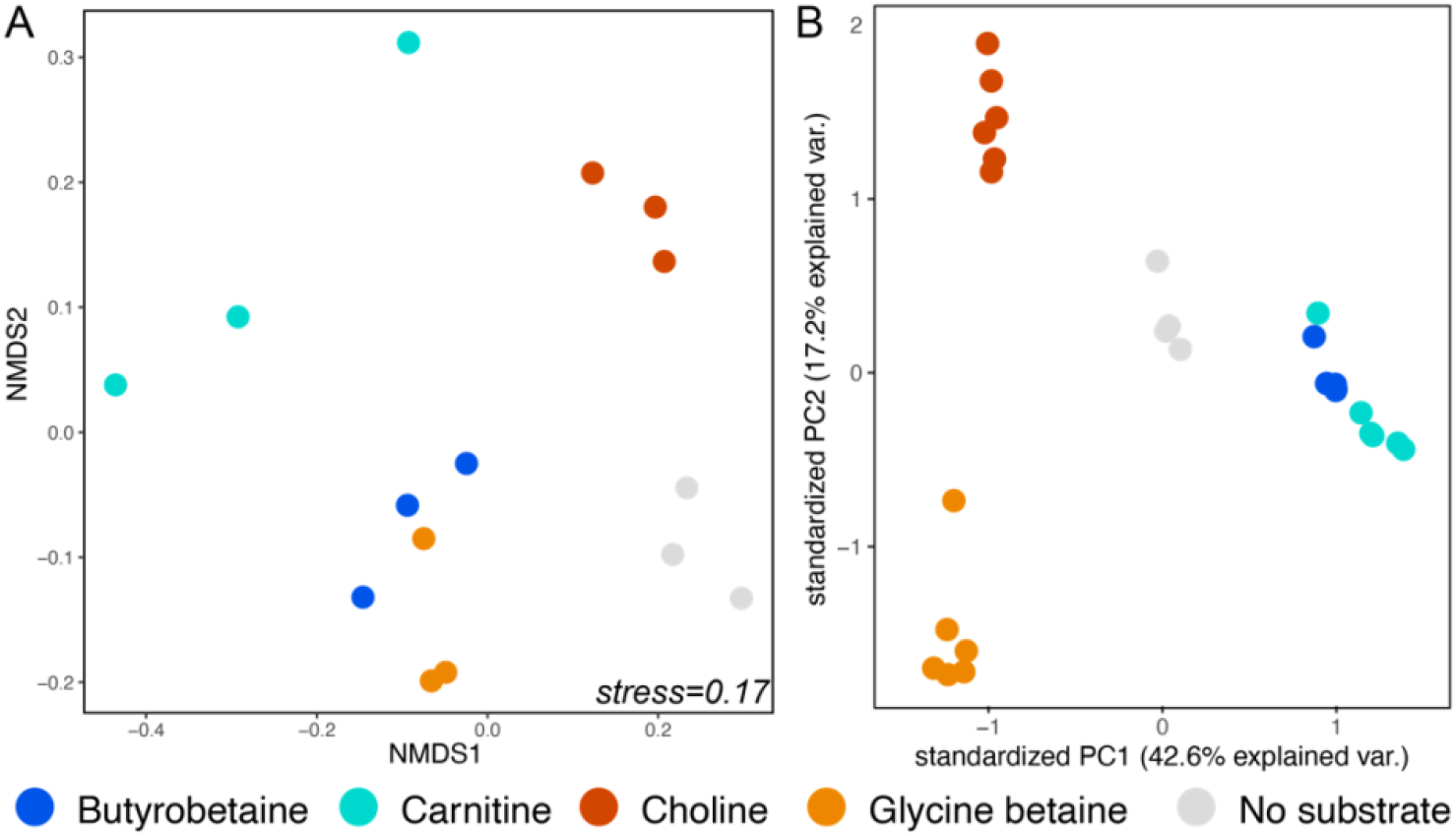
Ordinations show community wide gene expression (**A**, non-metric multidimensional scaling of final timepoints only) and metabolome (**B**, principal component analysis of T2 and TF timepoints) profiles for each reactor, with sample points colored by quaternary amine addition. Reactor microbial communities are statistically different by treatment and timepoint (mrpp, p<0.001); while 42% and 17.2% of variance in the metabolome is explained by the first and second components.

**Figure S5.**
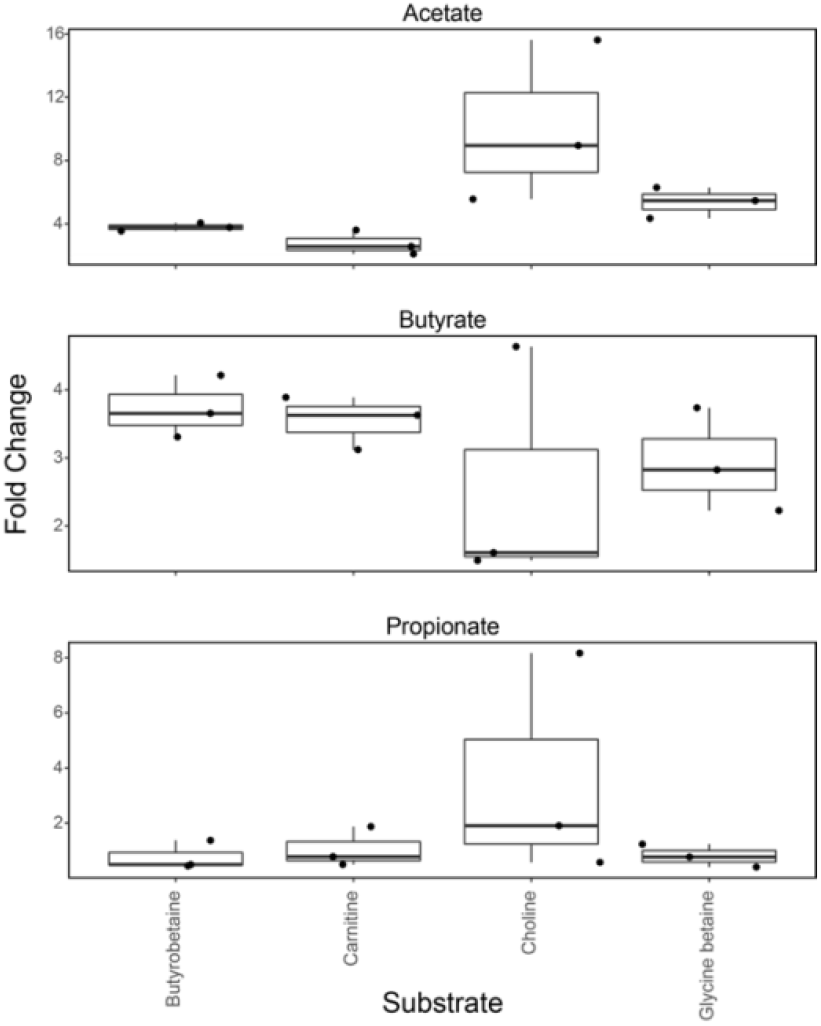
Fold change in SCFA concentrations detected in final timepoint of MA gut reactors relative to no substrate controls, with boxplots representing the triplicate reactor concentration for each reactor substrate. SCFA production gene expression is given in **Table S3**.

**Figure S6.**
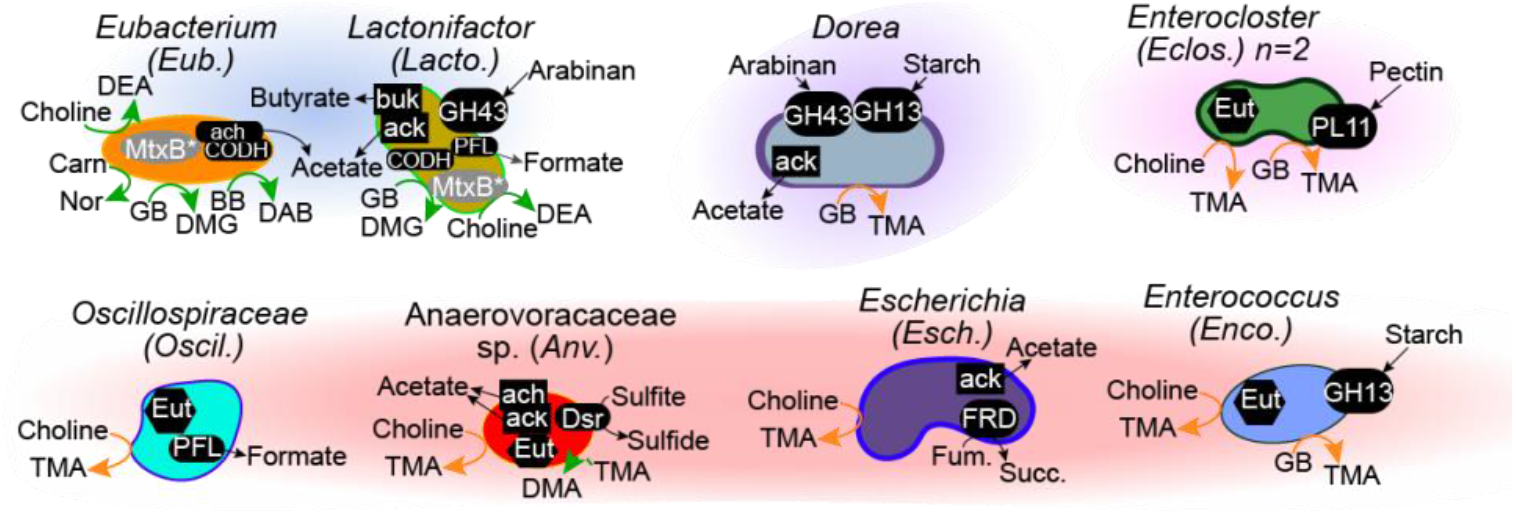
Illustrations inventory the genome-resolved expressed metabolism of the key MA metabolizing microorganisms. Green arrows denote nonTMA producing reactions, orange arrows denote TMA producing reactions, and black arrows denote non MA reactions that are relevant to gut homeostasis. These non MA genes include glycoside hydrolases (GH) for processing carbohydrates, as well as genes for SCFA production (*ack, buk, ach*) and respiration (*dsr, frd*). More detailed gene descriptions are provided in **Table S3**. Background cloud shading represents the conditions under which these microorganisms were inferred to be active. Blue cloud highlights *Eubacterium* and *Lactonifactor* which are responsible for subverting TMA concentrations across the quaternary amine stimulated fecal reactors. Purple cloud highlights the TMA producing *Dorea* exclusive to the glycine betaine reactor. The red cloud highlights three genomes from *Escherichia*, Anaerovoraceae, and Oscillospiraceae responsible for TMA production exclusive to the choline reactor. The pink cloud highlights the two genomes from the genus *Enterocloster* with the capacity to produce TMA from choline and glycine betaine.

**Figure S7.**
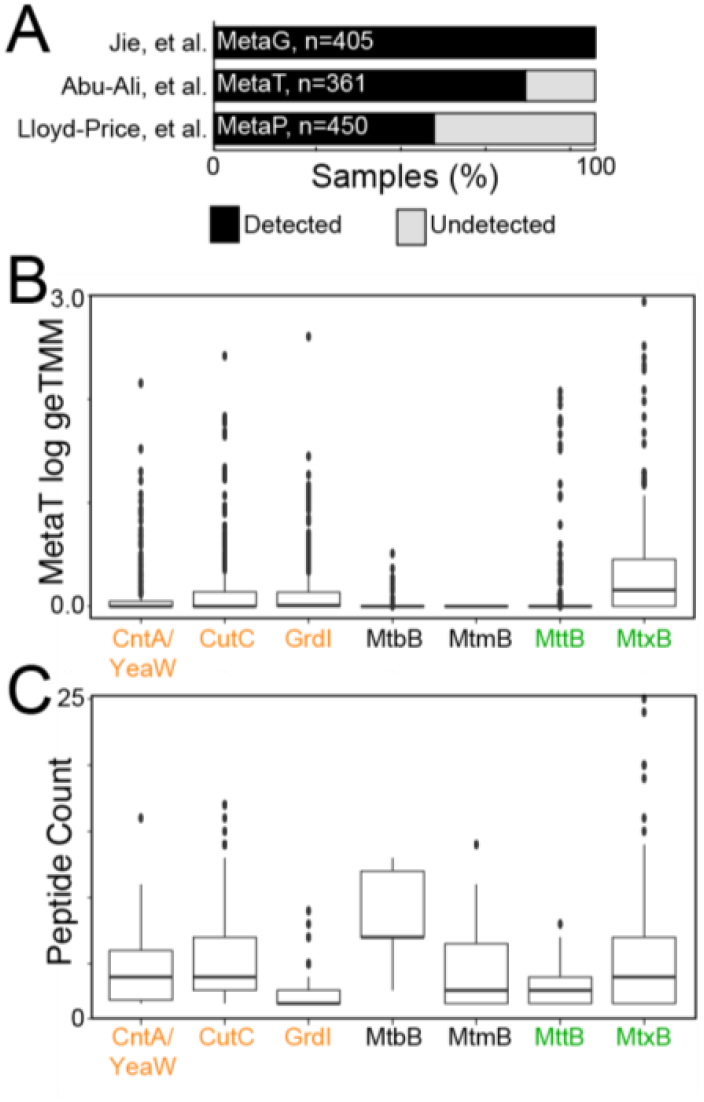
**A** Bar chart denotes the percentage of samples per study that members of MAGICdb are present or active. Studies include metagenomic data from a cohort of 218 individuals with atherosclerotic cardiovascular disease and 187 healthy controls, metatranscriptomic data from 361 adult men, and metaproteomic data from longitudinal sampling of 132 patients with irritable bowel syndrome. **B** Boxplots show the detection of MAGICdb genes in metatranscriptomics from 361 adult men from. Genes are colored by proatherogenic (orange) and nonTMA (green). **C** Boxplots show the detection of MAGICdb genes in metaproteomics from 132 patients with irritable bowel syndrome from ^66^. Genes are colored by proatherogenic (orange) and nonTMA (green). Across these studies, there was no significant difference in the number of expressed nonatherogenic and proatherogenic.

**Figure S8.**
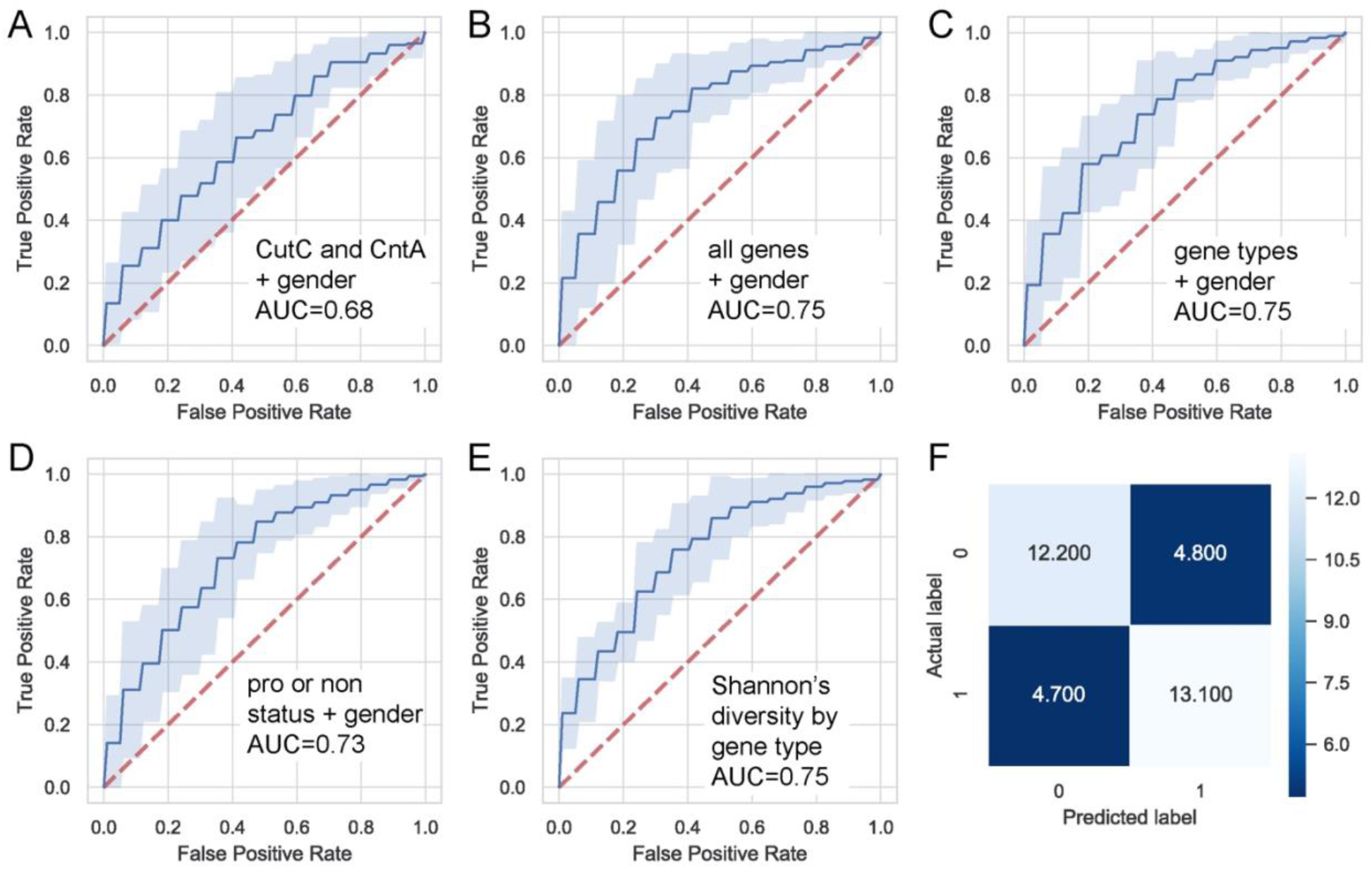
ROC curves of logistic regression models built using **A** abundance of *cutC, cntA/yeaW* summed per gene type + gender (genes used and model analysis similar to reported in ^32^, **B** abundance of all genes from MAGICdb + gender, each gene abundance in the unique MAGICdb gene database is included, **C** Abundance summed per gene type + gender, **D** Abundance of all genes summed per atherogenic status (proatherogenic and nonatherogenic) + gender, and **E** Shannon’s diversity of MAGICdb genes (no gender). **F** Confusion matrix for logistic regression model built based on the Shannon’s diversity of each type of gene in MAGICdb. Values are averaged over results from 10-fold cross validation.

